# Systematic analysis of human antibody response to ebolavirus glycoprotein reveals high prevalence of neutralizing public clonotypes

**DOI:** 10.1101/2022.01.12.476089

**Authors:** Elaine C. Chen, Pavlo Gilchuk, Seth J. Zost, Philipp A. Ilinykh, Elad Binshtein, Kai Huang, Luke Myers, Stefano Bonissone, Samuel Day, Chandrahaas R. Kona, Andrew Trivette, Joseph X. Reidy, Rachel E. Sutton, Christopher Gainza, Summer Monroig, Edgar Davidson, Erica Ollmann Saphire, Benjamin J. Doranz, Natalie Castellana, Alexander Bukreyev, Robert H. Carnahan, James E. Crowe

**Author notes:** Correspondence to: James E. Crowe, Jr., M.D., **Contact information: James E. Crowe, Jr., M.D.** *[LEAD CONTACT]*, Departments of Pediatrics, Pathology, Microbiology, and Immunology, and the Vanderbilt Vaccine Center, 11475 Medical Research Building IV 2213 Garland Avenue, Nashville, TN 37232-0417, USA.

## Abstract

Understanding the human antibody response to emerging viral pathogens is key to epidemic preparedness. As the size of the B cell response to a pathogenic virus protective antigen is undefined, we performed deep paired heavy and light chain sequencing in EBOV-GP specific memory B cells, allowing analysis of the ebolavirus-specific antibody repertoire both genetically and functionally. This approach facilitated investigation of the molecular and genetic basis for evolution of cross-reactive antibodies by elucidating germline-encoded properties of antibodies to EBOV and identification of the overlap between antibodies in the memory B-cell and serum repertoire. We identified 73 public clonotypes to EBOV, 20% of which encoded antibodies with neutralization activity and capacity to protect *in vivo*. This comprehensive analysis of the public and private antibody repertoire provides insight into the molecular basis of the humoral immune response to EBOV-GP, which informs vaccine design of new vaccines and improved therapeutics.

## Introduction

Periodic outbreaks caused by Ebola virus (EBOV) have occurred since its identification in 1976, when the virus caused two consecutive outbreaks in central Africa. Even in 2021, cases have occurred in Guinea and the Democratic Republic of Congo. EBOV causes severe or lethal disease in humans, with mortality rates ranging from 60 to 90%. Within the Ebolavirus genus, there are six species. Three of the six causes lethal disease in the humans, Zaire (EBOV), Bundibugyo (BDBV), Sudan (SUDV) (Feldmann et al., 2020). Taï Forest virus (TAFV) has caused one non-fatal case in humans, and Reston virus (RESTV) has caused asymptomatic infections in humans. Bombali ebolavirus, found by sequencing bats, can infect human cells but is not known to have caused illness in humans to date. Marburg virus (MARV), a closely related virus also in the Filoviridae family, also causes hemorrhagic fever with high mortality rates. These periodic outbreaks are of global health concern and highlight the need to accelerate Ebola virus disease (EVD) vaccines and therapeutics. Some EBOV-specific monoclonal antibodies (mAbs) neutralize virus and can mediate beneficial effects in humans with EVD (Bornholdt et al., 2019; Corti et al., 2016; Gilchuk et al., 2020b). And two mAb-based drugs Inmazeb^TM^ and Ebanga^TM^ received approval from the U.S. Food and Drug Administration (FDA) in 2020 (Levine, 2019; Pascal et al., 2018). Understanding the human antibody repertoire induced to ebolavirus GPs will help efforts to identify the next generation of pan-ebolavirus antibody therapeutics and aid design and development for broadly protective vaccines.

The genetic and functional diversity of the memory B-cell response and prevalence of public clonotypes to the EBOV GP remains unknown despite previous repertoire analysis of the B-cell response in individuals following vaccination with rVSV-ZEBOV (Ehrhardt et al., 2019) and following natural infection (Davis et al., 2019). Single-cell RNAseq methods now allow for isolation of authentically-paired heavy and light chain antibody variable genes, retaining the ability to functionally assay all B cells sequenced. This approach allows functional validation and profiling of antibodies at a large scale. The B-cell repertoire induced by EBOV vaccination or infection is likely to be diverse but has not been comprehensively characterized as a large data set from a single donor with paired heavy and light chain variable genes. Therefore, using single-cell paired heavy and light chain sequencing of the memory B cell response to EBOV GP allows us to (1) define the paired sequence repertoire and therefore accurate estimation of clonal diversity, (2) systematically select and characterize the functional diversity of repertoires, (3) understand evolution both on a genetic and functional basis, (4) identify antibodies shared in the memory B-cell repertoire and sera and the functions of those antibodies, and (5) quantify the prevalence and functionality of public clonotypes (clones seen in more than one individual).

Having a comprehensive in-depth understanding of the protective humoral response to EBOV GP on the repertoire level is important for devising optimal immunization schemes and informs the development of vaccines for multiple strains of Ebola (Cohen-Dvashi et al., 2020). In tandem, large-scale antibody studies could identify next-generation therapeutic antibody candidates. Such studies also can identify commonly induced antibodies that do not contribute to neutralization or protection, which is useful for building tools to benchmark the immunogenicity of new vaccine candidates. Understanding the genetic and functional diversity of the antibody repertoire to EBOV GP also can be therefore useful for vaccine development and therapeutic discovery.

Previous studies suggest that potent antibody response to GP early in convalescence is low, as potently neutralizing antibody responses appear later after recovery from infection (Davis et al., 2019; Williamson et al., 2019). This observation suggests that the neutralizing potency of antibodies to EBOV evolves through multiple rounds of affinity maturation during the process of somatic hypermutation. The antigenic landscape recognized by neutralizing antibodies also may evolve during convalescence. For instance, at early time points after recovery, most mAbs isolated target the glycan cap of GP, suggesting glycan-cap-specific antibodies may play a dominant role in the early human antibody response to EVD. A class of glycan-cap-specific antibodies are encoded by the *IGHV1-69* heavy chain gene, which specifies a germline-encoded complementarity-determining region 2 (CDRH2) with hydrophobic residues that facilitates binding to the glycan cap region (Murin et al., 2021). Therefore, germline-encoded *IGHV1-69*-antibodies likely play a role in the initial response to the GP. It has been shown that the functional profiles of several antibodies are retained when the somatically mutated sequences of such antibodies are reverted to the germline-encoded sequences (usually with minimal or no somatic mutations). Retention of function also has been reported for mAbs reverted in this way to germline-encoded sequences for other viral pathogens (Dong et al., 2021; Pappas et al., 2014; Yuan et al., 2020; Zhou et al., 2015; Zost et al., 2021). Identification of germline antibody genes encoding immunoglobulins with antiviral functional characteristics reveals a critical component of the early response to viral pathogens. Additionally, understanding how potent and cross-reactive antibodies evolve from germline-gene-encoded forms of those antibodies may inform rational vaccine design for the sites of vulnerability recognized by those mAbs (Rappuoli et al., 2016). Such studies, for example, define the critical residues and structures that should be retained at the site to induce antibodies with the desired functionality.

Humoral immunological memory is mediated in part by serum antibodies that are secreted by long-lived plasma cells, which usually live in the bone marrow. In contrast, memory B cells persist in circulation and are defined as long-lived and quiescent cells that are poised to quickly respond to antigen upon recall. Many antibody discovery efforts focus on memory B cells. Little is known about the composition of the polyclonal antibody secreted IgG repertoire and its overlap with the B cell receptors of memory B cells in EVD survivors. Defining the overlap of these two immune compartments could identify which clonal families contribute to the maintenance of protective humoral immunity.

Recent studies have suggested important overlap in individual’s antibody responses to infection and vaccination. Three classes of antibodies were described as encoded by the variable genes *VH3-15/VL1-40, IGHV3-13,* or *IGHV3-23* in multiple individuals following vaccination (Cohen-Dvashi et al., 2020) with the rVSV-ZEBOV (Ehrhardt et al., 2019) or ChAd3-ZEBOV(Rijal et al., 2019) vaccines, or natural infection (Bornholdt et al., 2016; Cagigi et al., 2018; Davis et al., 2019; Wec et al., 2017). Descriptions of these public clonotypes raises the important question of how much of the human B-cell response to EBOV is shared.

Mining the human repertoire for public clonotypes requires large numbers of antibody gene sequences, and validation of public clonotypes by testing recombinant immunoglobulins for specificity and antiviral function require antibody gene datasets containing authentically-paired heavy and light chain genes from single B cells. With larger paired-chain sequence datasets, the likelihood of identifying public clonotypes increases, allowing a functional understanding of the public antibody response to ebolavirus GPs. Mining for and understanding the properties of public clonotypes informs a deeper understanding of population immunity by revealing immunodominant B cell responses within immune populations, which may be of benefit for rational design of vaccines that may exhibit immunogenicity in a broader segment of the population. Knowing the public clonotype profile following natural infection also can enhance experimental vaccine testing, since the immunogenicity for desirable public antibodies recognizing cross-reactive sites of vulnerability for potent neutralization can be recognized at the cDNA sequence level.

To address this gap in knowledge, we sorted 100,000 EBOV GP-reactive memory B cells from a previously infected individual and performed large-scale single cell antibody gene sequencing. These sequences were used for in-depth analysis to define five points: (1) define and estimate the diversity of the paired sequence repertoire (2) characterize the functional diversity of repertoires, (3) understand evolution both on a genetic and functional basis, (4) identify antibodies shared in the memory B-cell repertoire and sera, and (5) quantify the prevalence and functionality of public clonotypes.

## RESULTS

### Identification of EBOV GP-specific memory B cells

To identify EBOV GP-specific memory B cells, we took PBMCs from a convalescent donor with history of EBOV infection in West Africa during the 2014 epidemic and performed a pan-B cell enrichment. Memory B-cells were then collected by flow cytometric sorting for IgM^-^ IgD^-^ CD19^+^ B-cells (**Figure 1A, B**). From this class-switched B cell population, we identified EBOV GP-reactive cells using biotinylated EBOV GP and fluorescently labeled streptavidin (**Figure 1B**). From 7 x 10^7^ total CD19+ B cells, we sorted ∼100,000 GP-reactive class switched B-cells; roughly 3% of the class-switched B cell population bound EBOV GP (**Figure 1C**). These GP-reactive cells then were single-cell sequenced using a single-cell encapsulation automated system (Chromium Controller; 10X Genomics). A total of 15,191 paired antibody heavy and light chain variable region sequences were obtained from the single-cell sequencing experiment. As a control experiment in parallel, a PBMC sample from a healthy adult with no history of exposure to EBOV also was subjected to the same workflow. For the control subject, IgM^-^ IgD^-^ CD19^+^ B-cells were sorted and subjected to single-cell sequencing to obtain 10,960 total paired heavy and light chain sequences for this non-specified-antigen-specific B cell set (**Figure 1C**).

**Figure 1.**
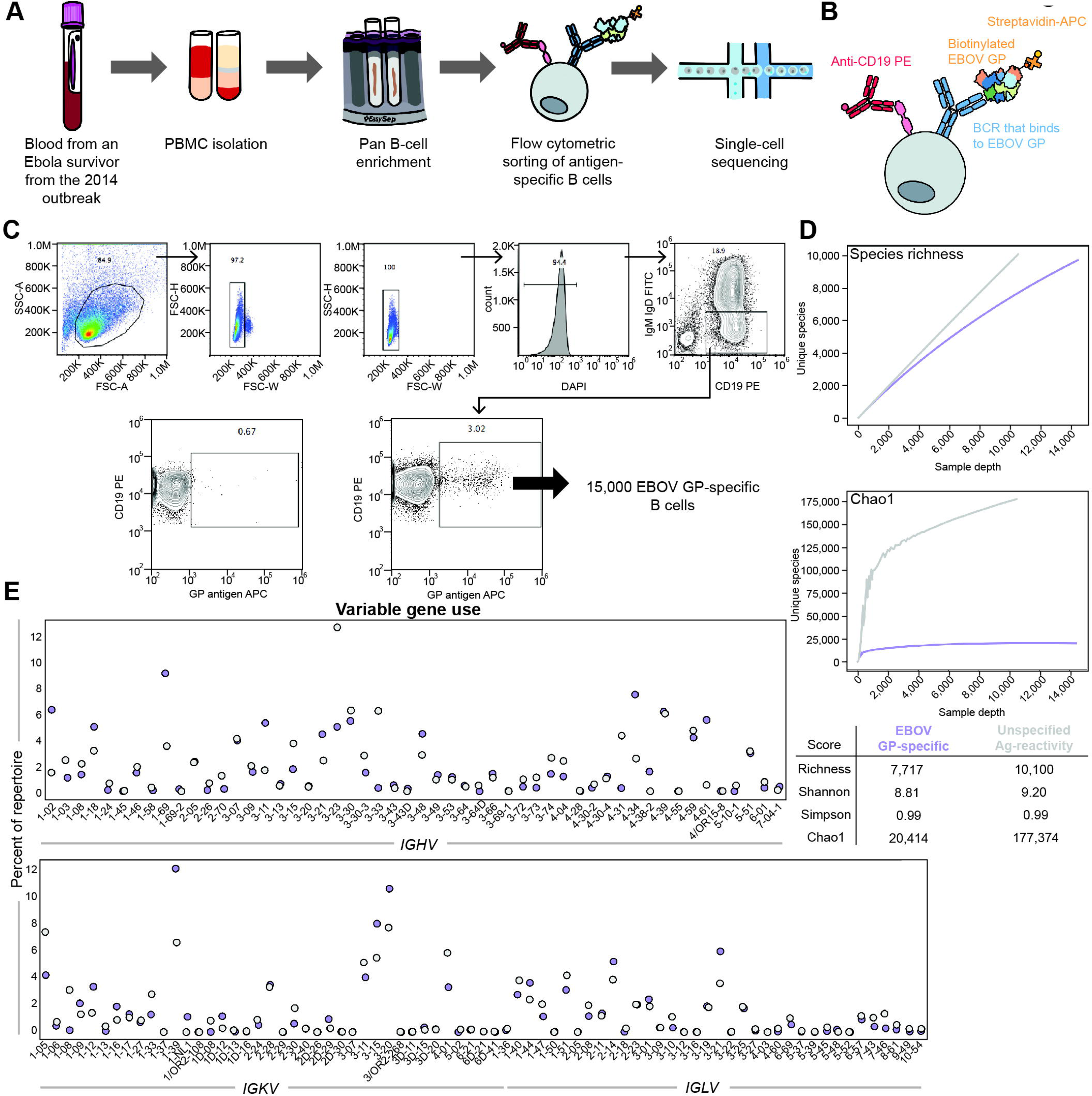
Identification of and diversity of EBOV GP specific memory B cells. **a.** Schematic of sample processing to identify and sort memory B cells. **b.** Schematic of flow cytometric staining to identify EBOV GP specific B cells. **c.** Gating for lymphocytes, singlets, live cells using DAPI, followed by class switched B cells. Cells were stained with anti-CD19 antibody conjugated to PE and anti-IgM and anti-IgD conjugated to FITC. EBOV GP was biotinylated and conjugated to streptavidin APC. FACS isolation of class-switched B cells (CD19^+^ IgM^-^ IgD^-^) specific to the EBOV GP (Antigen-APC) in a donor that has not previously been exposed to EBOV (left) or the convalescent donor (right) shown. **d.** Diversity metrics calculated for the EBOV GP-specific repertoire (purple) compared to the non-antigen-specific repertoire (grey). The first plot shows species richness and the second shows Chao diversity. The sample depth on the x-axis indicates the number of sequences, and the unique species on the y-axis indicates the number of clonal families. Additional diversity metrics were calculated including Shannon entropy and Simpson index. **e.** Variable gene usages in the heavy chain (top) or light chain (bottom) repertoire from sequencing. The number of sequences using each gene was calculated and normalized to a percentage using the total number of sequences as 100%. Purple dots indicate the EBOV GP-specific repertoire, grey dots indicate the non-antigen-specific repertoire.

### Genetic characteristics of the memory B cell repertoire to EBOV GP

Immunoglobulin features such as inferred variable gene use and percent identity to inferred germline genes were identified using the PyIR informatics pipeline based on IgBLAST (Soto et al., 2020; Ye et al., 2013). To examine the different variable genes used in antibodies specific to the EBOV GP that were captured from our GP-reactive B cell sorting experiment, the frequency of each *IGHV* or *IGKV/IGLV* gene used was measured and normalized to a percent using the total number of sequences acquired as 100%. The same analysis also was performed on the 10,960 total paired sequences for B cells obtained by from the control subject (**Figure 1C**). *IGHV1-69* was the most frequently used heavy chain variable gene (9.2%), followed by *IGHV1-02* (6.3%) and *IGHV4-34* (7.5%) (**Figure 1E**). For light chains, *IGKV1-39* was the most frequently used light chain variable gene (12.2%). Gene use for the non-specified-antigen-specific antibodies from the control subject are plotted for comparison. The median amino acid lengths of CDRH3 or CDRL3 for the EBOV GP-specific repertoire was 17 or 9, respectively. In comparison, the median amino acid lengths of CDRH3 or CDRL3 for the non-specified-antigen-specific repertoire were 15 or 9, respectively. For the EBOV GP-specific repertoire, the average identity to germline was 94.2% for the heavy chain and 96.1% for the light chain, with the median number of mutations being 16 or 10 amino acids respectively. As a comparison, in the non-specified-antigen-specific repertoire, the average identity to germline was 95.6% or 97.2% on the heavy or light chain, with the median number of mutations being 12 or 7 amino acids, respectively. These findings indicate that the EBOV GP-specific repertoire is slightly more mutated than the non-specified-antigen repertoire, with slightly longer CDR3s in heavy and light chains, although it is also possible that it is due to differences between individuals.

### Identification of clonal families

To identify clonal families to which each sequence belongs, sequences were clustered by binning the clones based on the inferred immunoglobulin heavy variable (*IGHV*) gene, immunoglobulin heavy joining (*IGHJ*) gene, and the amino acid length of the CDRH3. Then, sequences were clustered according to 80% nucleotide sequence identity in the DNA sequence encoding the CDRH3. Next, sequences were binned further based on the inferred immunoglobulin light variable gene (*IGLV* or *IGKV*) and immunoglobulin light joining (*IGLJ* and *IGKJ*) genes and 80% nucleotide sequence identity in the DNA sequence encoding the CDRL3. From the 15,191 total paired sequences derived from our EBOV GP-specific sort, 10,087 clonal families were identified. Of these, 6,923 were singlets (meaning there were no other sequences that clustered with that single sequence). Additionally, 2,382 were doublets, meaning two sequences clustered together, but did not cluster with any other sequence. We defined clusters as clonally expanded families if they included five or more sequences and found 224 such clonal families. To compare the distribution of EBOV GP-reactive clonal families to those of the control subject, we applied the same clustering scheme to the non-antigen-specific sequence set. From the 10,960 sequences in that individual, 10,527 clonal families were identified. From that set, 10,172 were singlets and 305 were doublets. Only 21 clonally expanded clonal families were observed in the control subject (**Figure 2B**).

**Figure 2.**
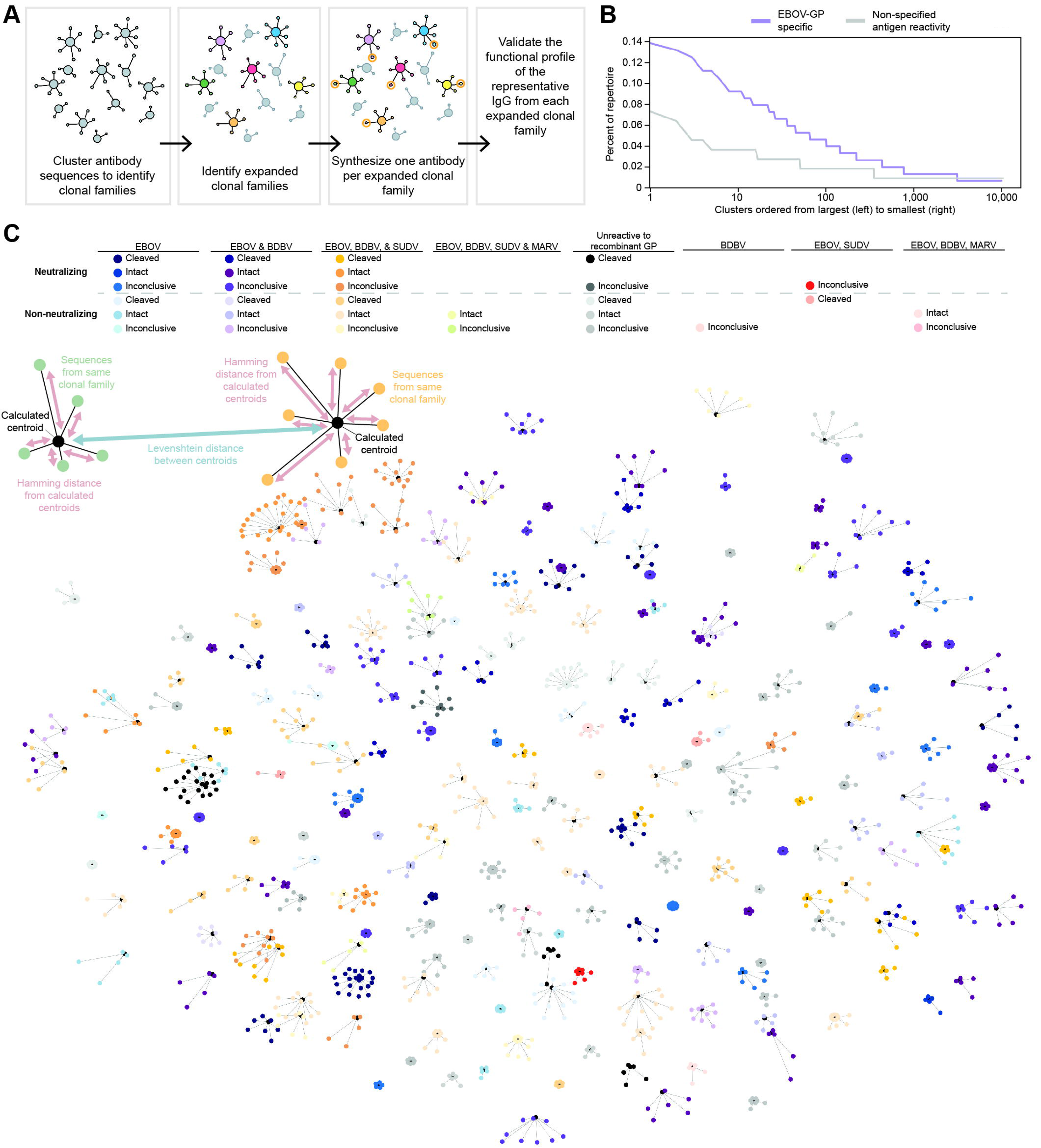
Analysis of the clonally expanded EBOV GP-specific repertoire. a. Schematic of identification of clonally-expanded families and selection of one clone per family. b. Graph showing the distribution of clonal families. After clustering was completed, clusters were ordered from the largest cluster to smallest cluster, then plotted in that order as the percent of the EBOV GP-specific repertoire (purple) and non-antigen-specific repertoire (grey). c. Network plot of the clonally-expanded repertoire with the functional characteristics of each clonal family plotted on and schematic showing how calculations were derived to construct network diagram. Reactivity to each glycoprotein in ELISA is denoted by different colors. Blue indicates antibodies monospecific to EBOV, purple indicates antibodies specific to EBOV and BDBV, orange indicates antibodies specific to EBOV, BDBV, and SUDV. Green indicates antibodies specific to EBOV, BDBV, SUDV, and MARV. Grey antibodies did not react with any GP tested in ELISA. Salmon indicates antibodies specific to BDBV only. Red indicates specificity for EBOV and SUDV. Pink indicates specificity for EBOV, BDBV, and MARV. Different shades of each color indicate neutralizing capacity, with the darker dots indicating neutralizing antibodies for VSV-EBOV and the lighter dots indicating non-neutralizing antibodies. Different shades within each color represent whether the antibody preferably bound to the intact GP (GP_ecto_) or cleaved GP (GP_cl_), or if it bound well to both.

We next estimated the size and diversity of the EBOV GP-reactive memory B cell repertoire in the convalescent donor using rarefaction analysis(Saary et al., 2017). In this work, we used the clonal families determined through the clustering scheme described above as taxonomic units or species. We first plotted species richness that is present in the defined sample set; species richness measures the number of species, or in our case clonal families. The species richness curve continually increased but never plateaued, meaning that as our sequencing depth increased, we continued to discover new clonal families. This finding suggested that even at this substantial depth of sequencing, we did not identify all EBOV GP-specific clonal families present in this donor. The same approach was used for the sequences from the control subject, for whom the species richness curve exhibited a steeper slope than the EBOV GP-specific curve. This comparison indicates the EBOV GP-specific sequence set is less diverse than the non-GP-specific sequence set that was captured in our experiments (**Figure 1D**).

Next, we calculated the Chao diversity index (Chao1) (Hsieh et al., 2016), which estimates the number of species there is likely to be in the sample set. The Chao diversity index for the EBOV GP-reactive antibody sequences at a sample depth of 90% of the repertoire was 20,329. This value suggests that there are an estimated 20,329 clonal families present in the EBOV GP-specific antibody repertoire within this donor, of which we identified 10,087 (∼50% of the total). The estimated total of the non-antigen-specific antibody sequence set at a sample depth of 90% is 177,374, from which we have identified 10,527 clonal families. However, the number for the non-GP-specific repertoire is not a confident estimate as this value grows at every new sample depth, and it is likely that the number of clonotypes in a non-GP-specific repertoire is much higher than estimated here. In contrast, the estimated number of clonal families stayed consistent with increasing depth for the EBOV GP-reactive antibody sequence set estimate, giving confidence in the estimated number of total species. This outcome can be visualized in **Figure 1D**, as the EBOV GP-specific repertoire unique species plot exhibits plateauing at a much lower sample depth. With more than a ten-fold difference in estimated diversity, this finding indicates that the EBOV GP-reactive antibody repertoire is much less diverse and has fewer clonal families compared to a GP-antigen-specific repertoire. Therefore, through modeling, we predicted that we have sampled about half of the EBOV GP-specific memory B-cell repertoire in the convalescent donor, as we have identified 10,087 clonal families out of the 20,329 estimated.

### Functional characterization of clonally expanded EBOV GP-specific repertoire

To understand the functionality of each of the 224 clonally-expanded families from the EBOV immune subject, the most somatically mutated member of each family was selected for antibody gene cDNA synthesis and recombinant IgG expression (**Figure 2A**), using previously described microscale production and purification approaches (Gilchuk et al., 2020a). Purified mAbs then were tested for binding in ELISA to recombinant trimeric EBOV GP, BDBV GP, SUDV GP, or MARV GP ΔTM proteins. Next, they were tested for binding to cleaved or intact GP displayed on Jurkat cells using a flow cytometric assay (Gilchuk et al., 2020b). The full-length membrane-bound EBOV GP molecules expressed on the surface of Jurkat cells likely are similar to the native form of GP on the surface of a viral particle or on naturally-infected cells (Davis et al., 2019). Lastly, neutralizing activity of the antibodies was assessed using real-time cell analysis (RTCA) assay, allowing for quantification of cytopathic effect induced by replication-competent VSV-EBOV (VSV-encoding EBOV GP in place of the VSV G protein)(Gilchuk et al., 2020a; Zost et al., 2020). The tests showed that 80% of mAbs recognized EBOV GP, 60% recognized BDBV GP, 30% recognized SUDV GP, and 2% recognized MARV GP. 11% of mAbs recognized the three lethal strains of ebolavirus: EBOV, BDBV, and SUDV. Also, 30% of mAbs preferentially bound to cleaved GP, while 34% of mAbs preferentially bound to intact GP. Lastly, 95 mAbs neutralized VSV-EBOV (**Figure S1**). In conclusion, 80% of the clonally-expanded repertoire selected reacted with EBOV GP, with 42% showing neutralizing properties to VSV-EBOV.

### Building a network of the clonally expanded population

As there are multiple antibodies within each clonal family/lineage, we sought to visualize the relationships of clones between and within lineages. We built a network combining the genetic similarities of antibodies within lineages, between different lineages, and the functional profile of each lineage through the experimental data determined from experiments described above.

To retain paired heavy and light chain sequence information, we selected matching CDRH3 and CDRL3 amino acid sequences and linked the two with an arbitrary string, allowing for the resulting single string to be used as a node, representing a single antibody. A centroid also was computed for each lineage of antibodies using Vsearch to represent the average CDRH3 and CDRL3 sequence for each clonal family. Hamming distances then were calculated from each of the linked antibody CDR3 sequences to the calculated centroids to investigate the relationships of antibodies within clonal families. As using Levenshtein distance accommodates for the different CDR3 lengths that can be present between different clonal families/lineages, the Levenshtein distances were calculated between each centroid representative of each clonal family to investigate the relationship between clonal families (**Figure 2C**). The functional characteristics of each clonal family (**Figure S1**) then were mapped onto the network, with all the clonal families colored by functional profile. Here, we can visualize the diversity of functional phenotypes within the clonally expanded repertoire (**Figure 2C**) in conjunction with genetic similarities of antibodies within and between clonal families revealing that there is a large set of neutralizing antibodies that preferentially bound to the cleaved GP, but the CDR3 similarities of these antibodies vary. This plot revealed a cluster of cross-reactive neutralizing antibodies with similar CDR3s on the upper left portion of the network plot (**Figure 2C**). Understanding relationships of such could be useful in predicting antibody function through sequence analysis.

### Overlap in the repertoire between B cell receptors encoded in the memory B cell population and immunoglobulins present in plasma

We recently described a proteo-genomic analysis for identifying EBOV-specific immunoglobulin proteins in convalescent human plasma from the same donor we used in this study (Gilchuk et al., 2021). EBOV GP-specific polyclonal antibodies from the donor plasma were purified and subjected to high-resolution liquid chromatography coupled to tandem mass-spectrometry, yielding sequences of antibody proteins present in plasma. Immunoglobulins in the plasma repertoire were identified as present in the memory B cell repertoire if there was over 50% coverage in the CDR3 region and a general peptide coverage over 100% of the CDR3, as previously described (Gilchuk et al., 2021). A subset of 1,512 EBOV GP-specific memory B cell antibody variable gene sequences was used for the original study. Here, since we had obtained a much (10-fold) larger memory B cell repertoire from heavy and light chain paired sequences from this individual, we reinvestigated the portion of antibodies that is shared between the plasma immunoglobulin protein and memory B cell receptor repertoires.

Despite the 10-fold increase of gene sequences against which we could search, we only found an additional 82 antibodies, bringing the total antibodies found present in the overlap of antibodies in the plasma and in the memory B cell repertoire to 153. Of these, the clones belonged to 106 clonal families. A subset of 24 was from clonally-expanded clusters, and 44 were from singlets (**Figure 3A**). Of the 24 antibodies present in the plasma that came from clonally-expanded families, 17 of the antibodies neutralized VSV-EBOV. It had been previously described that polyclonal IgG isolated from convalescent plasma demonstrates preferential binding to cleaved GP(Davis et al., 2019; Gilchuk et al., 2021). However, of the 24 antibodies present in the plasma from clonally-expanded families, 16 of 24 bound preferentially to intact GP (**Figure 3B**). Therefore, it is likely that many of the polyclonal antibodies found in the plasma come from non-clonally-expanded memory B cell families. When identifying the V_H_ gene usages of these serum identified antibodies, the highest used genes were IGHV1-2 and IGHV3-11 at 10%. Following, IGHV4-34, IGHV3-21, IGHV1-69, and IGHV4-59 at 8%, 8% 7%, and 6% of the total serum antibodies identified respectively. Additionally, despite the large amount of gene sequences used here as a reference set for the proteomics studies, there appears to be a relatively small overlap between antibodies in the plasma and B cell receptors in the circulating memory B cell population.

**Figure 3.**
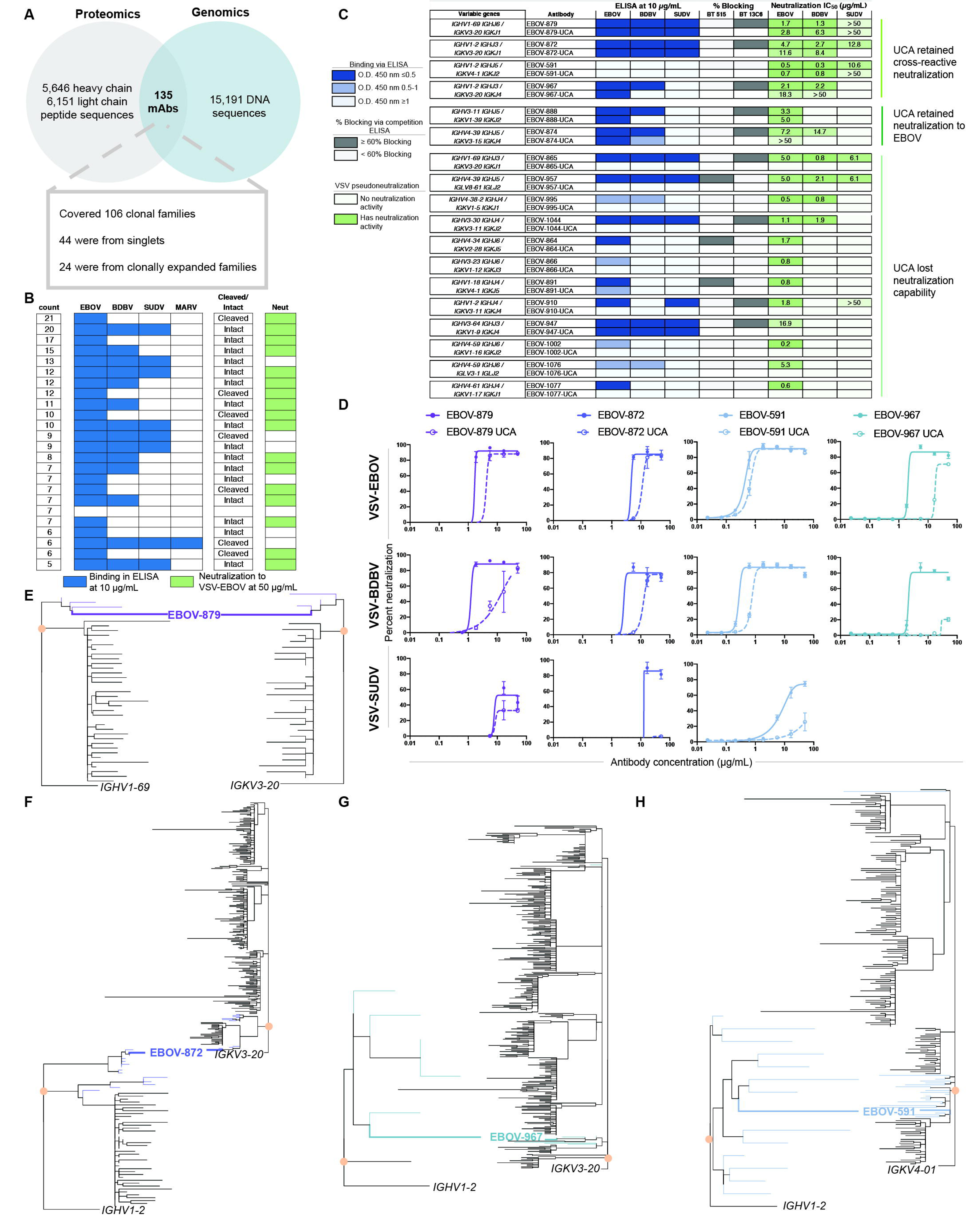
Characteristics of plasma antibodies and unmutated common ancestors of clonally-expanded antibodies. a. Venn diagram detailing the identification of antibodies present in both plasma and the memory B-cell repertoire. b. Characteristics of antibodies present in both the plasma and memory B-cell repertoire that originated from clonally-expanded families. The number of antibodies present in the clonal family is shown in the first column, followed by blue color indicating binding reactivity in ELISA to the different GPs, followed by binding to cleaved or intact GP, followed by neutralization for VSV-EBOV indicated in green. The empty box in the Cleaved/Intact preferential binding column indicates equivalent binding in that assay. Experiments were performed in biological duplicate, and the compilation of replicates is shown. c. Antibodies and their inferred UCAs with the functional profiles of each antibody. Gene use is listed in the first column followed by antibody names. Antibodies are listed with the mutated version of the antibody on the top row and the unmutated common ancestor (UCA) version on the bottom. The blue boxes indicate binding in ELISA to the different GPs at 10 µg/mL of antibody. The grey boxes indicate percent blocking in competition-binding ELISA against biotinylated EBOV-515, a base-region-specific reference antibody or against the glycan cap reference antibody 13C6. The green boxes indicate neutralization for VSV-EBOV, -BDBV, or -SUDV. The numbers inside the boxes indicate the calculated IC_50_ value for each antibody. Experiments were performed in biological duplicate and technical triplicates with similar results. A biological replicate from a single experiment is shown. d. Neutralization curves of unmutated common ancestor antibodies (dotted lines) that retained cross-reactive neutralization and their mutated counterparts (solid lines) against VSV-EBOV, -BDBV, or -SUDV. Experiments were performed in biological duplicate and technical triplicates with similar results. A biological replicate from a single experiment is shown. e. Maximum likelihood phylogenetic tree of the EBOV-879 lineage. The inferred UCA is indicated in the orange circle on the heavy chain and light chain tree. Blue lines indicate antibody sequences that were found in paired chain sequencing; black lines indicate sequences that were found in unpaired chain bulk sequencing that clustered with the clonal family. f. Maximum likelihood phylogenetic tree of the EBOV-872 lineage. The inferred UCA is indicated in the orange circle on the heavy chain and light chain tree. Blue lines indicate those antibodies found in paired sequencing; black lines are bulk sequences that clustered with the clonal family. g. Maximum likelihood phylogenetic tree of the EBOV-967 lineage. The inferred UCA is indicated in the orange circle on the heavy chain and light chain tree. Blue lines indicate those antibodies found in paired sequencing; black lines are bulk sequences that clustered with the clonal family. h. Maximum likelihood phylogenetic tree of the EBOV-591 lineage. The inferred UCA is indicated in the orange circle on the heavy chain and light chain tree. Blue lines indicate those antibodies found in paired sequencing; black lines are bulk sequences that clustered with the clonal family.

### Unmutated common ancestors of expanded clones reveal germline reactivity of clone encoded by *IGHV1-69* or *IGHV1-02*

Although we obtained unprecedented depth for paired heavy and light chain variable gene sequencing from single antigen-specific B cells from this donor, the depth of sequencing that can be acquired by bulk heavy or light chain antibody variable gene sequencing (without pairing) is still far superior. Therefore, to investigate the evolution of cross-reactive antibodies, we clustered sequences obtained from single cell paired sequencing with those obtained from bulk sequencing from the same leukapheresis sample PBMCs from this donor and constructed phylogenetic trees detailing the evolution of these cross-reactive antibodies versus that of monospecific antibodies.

Several clonally-expanded neutralizing antibodies with varying reactivities to the different GP and predicted epitopes were selected for further investigation of their evolution. To increase the amount of sequencing to use for phylogenetic analysis, heavy and light chain bulk sequencing was performed on PBMCs originating from this donor without antigen-specific sorting. The heavy chains from all clonal families previously identified in the single-cell sequencing were then clustered with those in the heavy chain bulk sequencing, and the light chains were clustered similarly. Sequences were clustered based on the same V and J gene usage as well as 80% identity of the CDR3 nucleotide sequences. Using the clustered sequences for each clonal family, maximum likelihood trees were constructed for both heavy and light chains, and the unmutated common ancestor (UCA) was inferred for each clonal family.

To determine whether the antiviral function of each clonal family was due to germline-encoded reactivity or due to somatic mutations that accumulated during antibody evolution, we investigated the binding and neutralization profile for each inferred UCA (**Table S1**). Each UCA antibody was expressed and tested for binding to EBOV, BDBV, or SUDV GP, and neutralization against VSV-encoding EBOV GP, BDBV GP, or SUDV GP in place of the VSV G protein (VSV-EBOV, VSV-BDBV, or VSV-SUDV respectively). Of the 18 UCAs tested, 12 lost their ability to bind to the different GPs in comparison to their mutated counterparts (**Figure 3C**).

Two UCA antibodies not only still bound to the appropriate GPs, but also they maintained their ability to neutralize VSV-EBOV, albeit with lower potency. EBOV-888-UCA uses *IGHV3-11*/*IGKV1-39*. EBOV-874-UCA uses *IGHV4-39*/*IGKV3-15* and maintained capacity to neutralize VSV-EBOV with reduced potency but lost its ability to neutralize VSV-BDBV. Therefore, it is likely that these two gene combinations contribute to germline-encoded neutralization properties specific to EBOV.

Additionally, four UCA antibodies retained the ability to mediate cross-reactive neutralization: EBOV-879-UCA, EBOV-872-UCA, EBOV-591-UCA, and EBOV-967-UCA. EBOV-879-UCA is encoded by *IGHV1-69/IGKV3-20* and neutralized VSV-EBOV, -BDBV, and -SUDV. These data show that antibodies encoded by *IGHV1-69/IGKV3-20* possess germline-encoded capacity to neutralize across all three medically important ebolavirus species, and they acquired increased potency during the process of somatic hypermutation (**Figure 3D,E**). EBOV-872-UCA uses *IGHV1-2/IGKV3-20* and maintained neutralization against VSV-EBOV and VSV-BDBV. As EBOV-872-UCA lost its ability to neutralize VSV-SUDV, it is likely that EBOV-872 acquired the capacity to neutralize SUDV by acquiring somatic mutations (**Figure 3D,F**). EBOV-967-UCA also maintained its cross-reactive neutralizing activity for both EBOV and BDBV, even though its neutralization potency for BDBV is relatively low at 50 µg/mL (**Figure 3D**). Therefore, it is likely that these antibodies started off with neutralizing properties mostly for EBOV with weak inhibition of BDBV but evolved to gain potency for the two strains (**Figure 3G**). EBOV-591-UCA also retained its ability to neutralize all three strains, however, its potency to SUDV dropped substantially. EBOV-591 also is encoded by *IGHV1-2* but uses a different light chain gene, *IGKV4-1* (**Figure 3D, H**).

EBOV-967 uses *IGHV1-2/IGKV3-20*, the same V_H_ and J_H_ genes as EBOV-872, however they differ in their J_L_ gene usage. As both antibodies neutralized EBOV and BDBV, it is likely that *IGHV1-2/IGHJ3* in combination with *IGKV3-20* encodes for neutralization of EBOV and BDBV (**Figure 3C, D**). We note all UCA antibodies that neutralized virus targeted the glycan cap region of the GP, since they competed for binding with the glycan cap antibody 13C6. These results indicate that germline-encoded structural features contribute to the ability of these antibodies to neutralize virus (**Figure 3C**).

### 73 public clonotypes are identified

We curated a database containing Ebola-specific mAbs by combining the large set of sequences acquired here with several smaller sets of previously reported Ebola-specific antibodies (Davis et al., 2019; Ehrhardt et al., 2019; Rijal et al., 2019; Wec et al., 2017). Collectively, this database includes sequences from 12 individuals determined following either natural infection or vaccination. These sequences then were clustered to identify public clonotypes. Sequences were first binned by their V and J gene use and CDR3 length. Next, sequences were clustered by 60% on the CDR3 nucleotide sequence and binned by the light chain V and J gene. Clusters with sequences from two or more of the 12 individuals then were identified as public clonotypes. A total of 73 public clonotypes were identified. One public clonotype was shared among 6 donors. Another was shared among four donors. Five public clonotypes were shared between three donors, and the remaining were all shared between two donors (**Figure 4A**). All 294 members of the 73 public clonotypes were synthesized and expressed as recombinant IgGs as previously described (Gilchuk et al., 2020a) and tested by ELISA for binding to EBOV, BDBV, SUDV, MARV GP, or EBOV sGP. Next, they were tested for binding to cleaved EBOV GP (GP_cl_) or intact EBOV GP (GP_ecto_), and for neutralization of VSV-EBOV. As most members of each public clonotype were expected to share similar functional profiles due to genetic similarity, the predicted functional profile was determined by identifying the dominant functional phenotype in each public clonotype (**Figure 4C, S5**).

**Figure 4.**
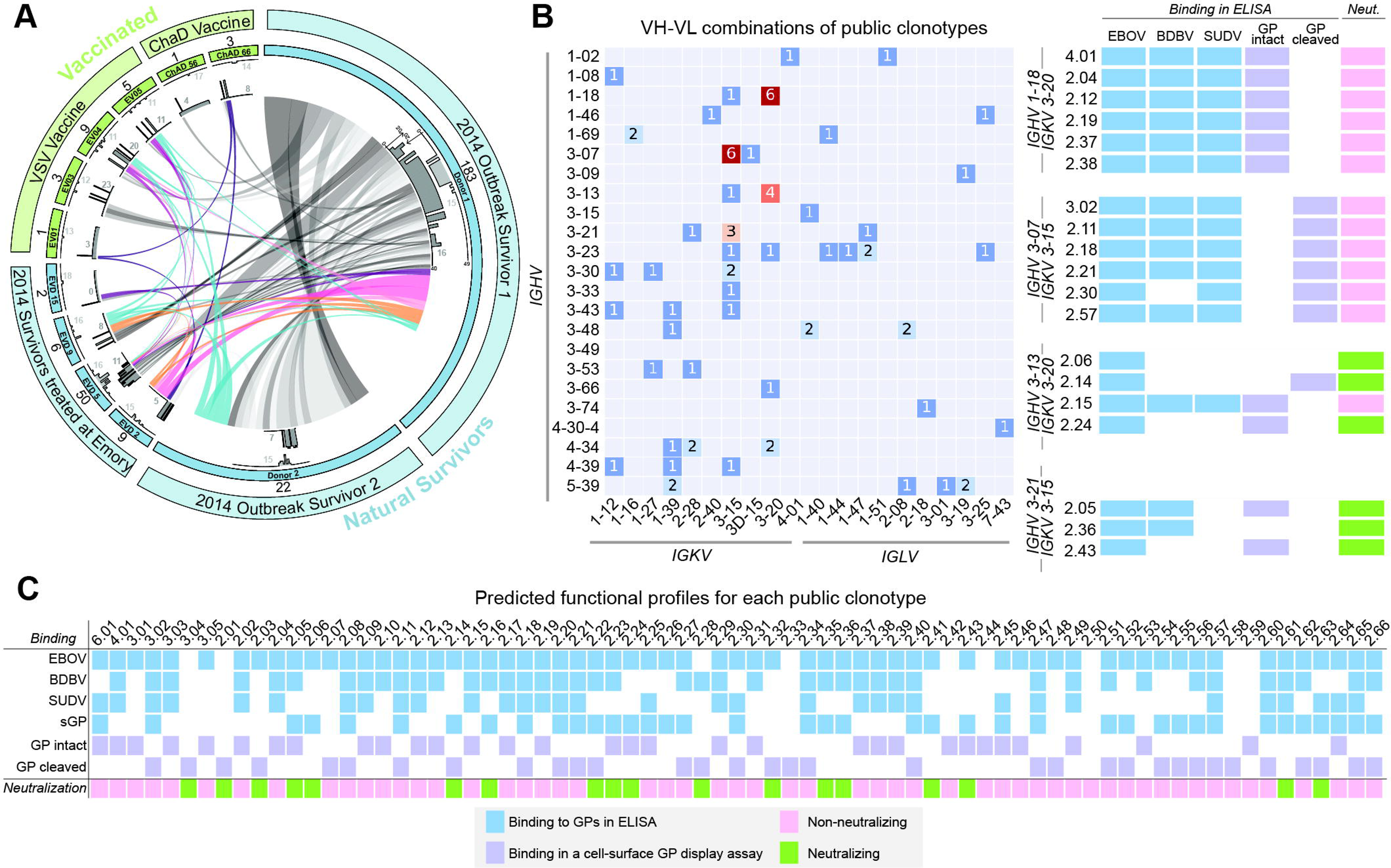
Identification of public clonotypes. a. Clonal overlap between each vaccinated (green) and convalescent (blue) donors. Numbers inside the first outer circle indicate the number of sequences that were identified as public clonotypes from the respective donor. The light grey color shows the distribution and median CDR3 length. The dark grey color shows the distribution and median number of somatic mutations of the public antibodies from that donor. b. Heavy and light chain variable gene usage combinations for all public clonotypes identified. Numbers inside boxes indicate the number of public clonotypes using that gene combination. Public clonotype groups using highly used genes are listed on the right. Blue indicates binding to GPs in ELISA, purple indicates binding via cell-surface GP display assay; pink indicates non-neutralizing, and green indicates neutralizing. c. Functional profiles of each of the 73 public clonotypes after expression and functional testing of the 294 public clonotype antibodies and inferring a functional profile for each of the public clonotype groups. Blue indicates binding to GPs in ELISA, purple indicates binding in a cell-surface GP display assay; pink indicates non-neutralizing, and green indicates neutralizing. Experiments were performed in biological duplicates. A compilation of the average of all experiments is shown.

From the 73 public clonotypes, there was a diversity of variable heavy and light chain combinations used. However, the two most frequent combinations observed were *IGHV1-18/ IGKV3-20* and *IGHV3-07/IGKV3-15.* All public clonotypes that used *IGHV1-18/IGKV3-20* bound to EBOV, BDBV, and SUDV GP, but none neutralized VSV-EBOV. All but one of the public clonotypes that used *IGHV3-07/IGKV3-15* bound to EBOV, BDBV, and SUDV GP; the outlier bound to EBOV and SUDV but not BDBV GP. None of these public clonotypes exhibited neutralization to VSV-EBOV. Therefore, it is likely that GP-reactive antibodies reacting to EBOV, BDBV, and SUDV using *IGHV1-18/IGKV3-15* and *IGHV 3-07/IGKV3-15* are found in many individuals. Additionally, there were four public clonotypes that used *IGHV3-13/IGKV3-20*, and three that used *IGHV3-21/IGKV3-15*. The majority of these seven public clonotypes had neutralizing properties. Additionally, the bulk of the public clonotypes using these variable genes had similar functional profiles, hinting that these combinations of variable genes may encode the neutralization properties for VSV-EBOV (**Figure 4B**).

### 15 of 73 public clonotypes neutralize EBOV

Of the 73 public clonotypes, 15 neutralized VSV-EBOV GP. Members of these 15 public clonotypes then were tested for binding to EBOV, BDBV, and SUDV GP. One of the 15 public clonotypes bound to all three GPs, two of 15 bound to EBOV and BDBV GP, nine of 15 bound to only EBOV GP, and three of 15 did not exhibit binding to any GPs, indicating that the majority of the neutralizing public antibody response is primarily monospecific (**Figure 5A**).

**Figure 5.**
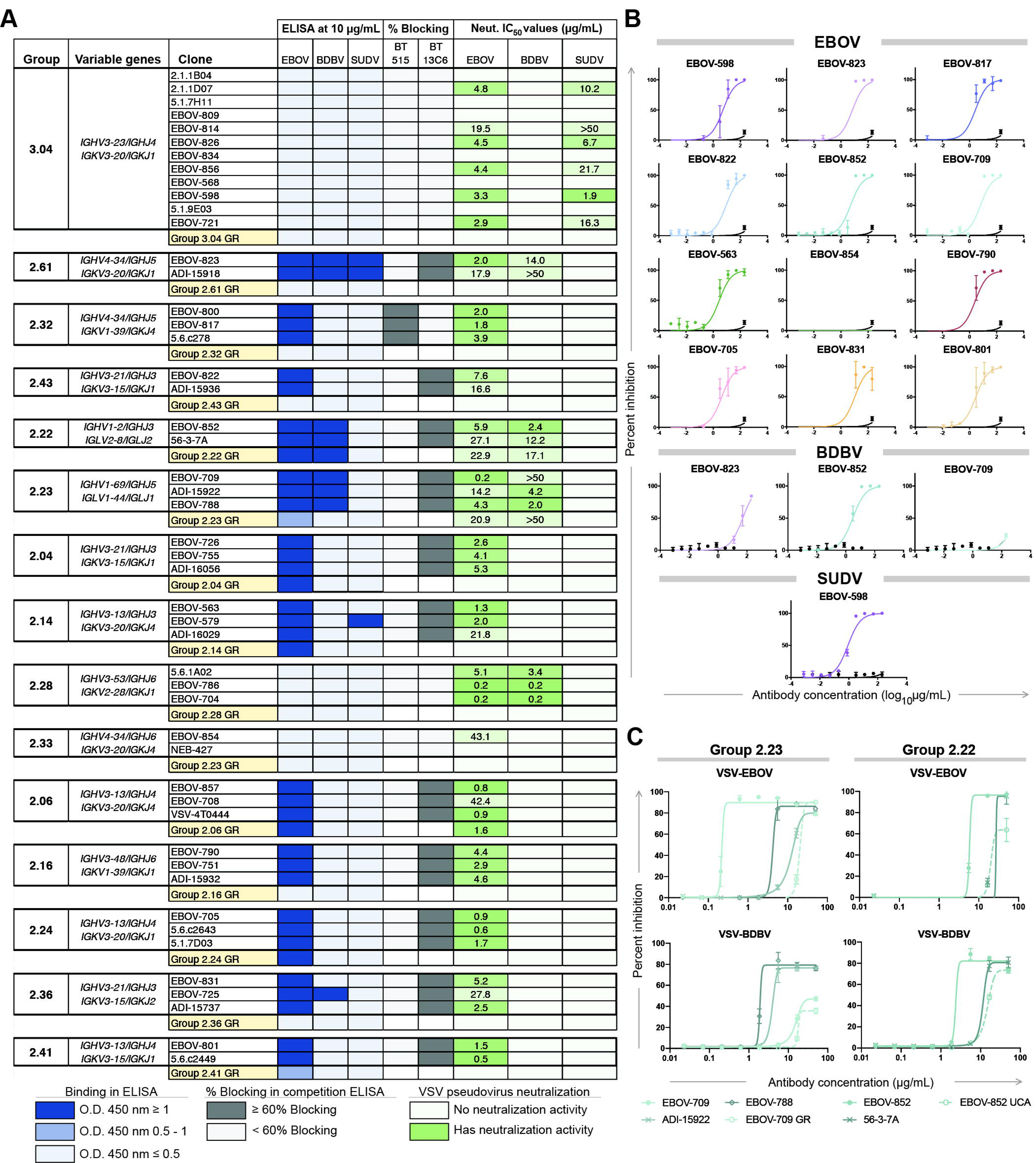
Properties of neutralizing public clonotypes. a. Table showing all 15 neutralizing public clonotypes. The first column identifies the public clonotype group number, the second column details the variable gene usage. The third column indicates clone name, with all the public clonotype antibodies in the group indicated in white and the germline revertant version of that group’s antibody in yellow. Blue boxes indicate binding in ELISA at an antibody concentration of 10 µg/mL. Grey indicates percent blocking in a competition-binding assay. Green indicates neutralization for VSV-EBOV, -BDBV, or -SUDV with the IC_50_ values written inside the boxes. Clone names highlighted in yellow at the bottom row of each section is the germline revertant version of that public clonotype and its respective functionality. b. Authentic virus neutralization curves for a representative antibody of each public clonotype group. c. Neutralization curves from antibodies that retained cross-reactive neutralization at the germline level. Group 2.23 antibodies to VSV-EBOV and -BDBV are shown on the left and Group 2.22 antibodies to VSV-EBOV and -BDBV are on the right. Dotted lines in each graph indicate the germline revertant antibody curve. Solid lines in varying colors indicate the matured versions of the antibodies in that public clonotype group.

Next, all mAbs were tested for neutralization of VSV-EBOV, -BDBV, or -SUDV. Although most public clonotypes only exhibited neutralization to VSV-EBOV, Group 3.04 neutralized both VSV-EBOV and -SUDV. Groups 2.61, 2.22, 2.23, and 2.28 neutralized VSV-EBOV and - BDBV (**Figure 5A**).

One mAb from each public clonotype group then was tested for neutralization of authentic virus. The neutralization profile for each mAb previously established with VSV-EBOV was reflected in the authentic virus neutralization assay except for EBOV-854. EBOV-854 exhibited a low neutralization potency in the VSV neutralization experiment and did not show any neutralization in the authentic virus experiment. Together, these findings verified that we identified public clonotypes exhibiting neutralization properties for EBOV, BDBV, and SUDV (**Figure 5B**).

### Most of the neutralizing public clonotypes identified target the glycan cap

All mAbs within the public clonotype groups were competed against each other for binding to EBOV GP in a competition-binding ELISA for pairwise comparison (**Figure 6A**). As this pairwise competition-binding ELISA was done with intact IgG, it is likely that the flexibility of the Fc region of the mAbs resulted in the asymmetric competition-binding grid in several public clonotype groups. Despite asymmetric competition (**Figure 6A**), all mAbs within each public clonotype group competed against each other. All mAbs then were tested for competition-binding with the previously epitope-mapped mAbs EBOV-515 (a base antibody) or 13C6 (a glycan cap antibody). Of the 15 neutralizing public clonotypes, 11 targeted the glycan cap and 1 targeted the base region of GP (**Figure 5A**) as concluded from competition-binding ELISA results. The remaining three antibodies did not bind to GP in ELISA and therefore, we used negative stain electron microscopy (EM) to identify the antigenic site recognized by these neutralizing antibodies.

**Figure 6.**
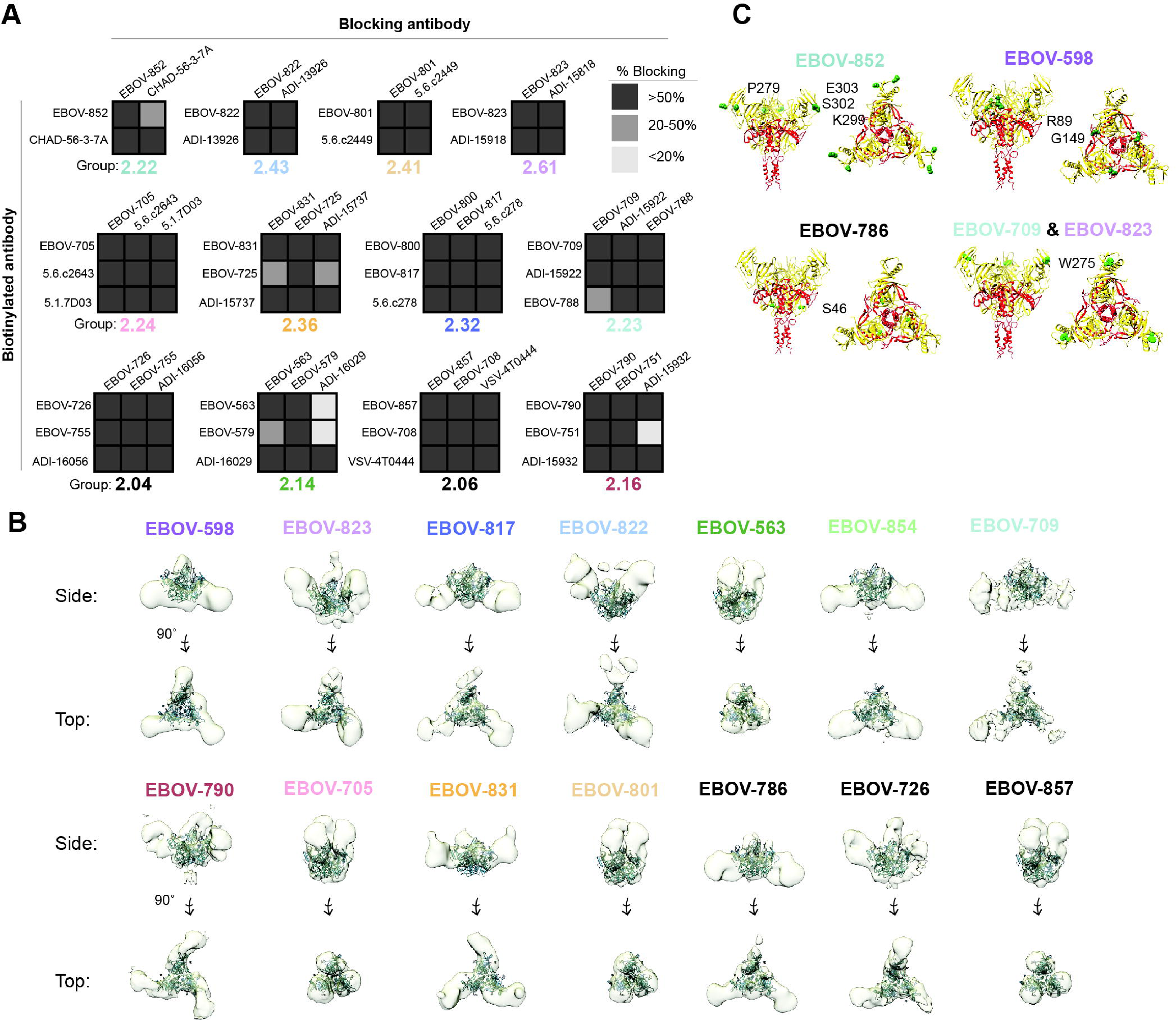
**Epitopes targeted by neutralizing public clonotypes.** a. Competition-binding ELISA results for antibodies within each public clonotype group in competition with each other. Unlabeled blocking antibodies applied to the GP antigen first are listed across the top of each grid while the biotinylated antibodies that are added to the antigen-coated wells second are indicated on the left. The number in each box represents the percent un-competed binding of the biotinylated antibody in the presence of the indicated competed antibody. The experiment was performed in biological duplicate and technical triplicates with similar results. A biological replicate from a single experiment is shown. b. Negative stain EM of EBOV GP in complex with Fab forms of different antibodies. 3D reconstructions are shown. The Fab (blue) is docked to a trimer of the EBOV GP (grey). c. Critical binding residues for EBOV-852, EBOV-598, EBOV-786, EBOV-709, and EBOV-823 as determined by loss of binding in alanine-scanning GP mutagenesis studies. Residues where alanine caused loss of binding for antibodies are indicated in green.

Fab-EBOV GP complexes were imaged with a representative mAb from each public clonotype group (**Figure 6B, S4**). Although EBOV-598 (Group 3.04), EBOV-786 (Group 2.28), and EBOV-854 (Group 2.33) and other members of their respective public clonotype groups did not exhibit binding to GP in ELISA they did show binding to GP_cl_ in a cell-surface display assay and had neutralization properties (**Figure 4C, 5A, S5**); these mAbs complexed with EBOV GP for EM studies, and 3D reconstructions were made. These low-resolution reconstructions show that all three of these mAbs as well as EBOV-817, which competed for binding with the reference antibody EBOV-515, bind the base region of GP. Although EBOV-852 was visualized on the grid, and we were able to obtain 2D images of it in complex with GP, we were unable to obtain a 3D reconstruction for it. Low-resolution reconstructions of the rest of the public clonotypes show that the public clonotypes bind diverse regions of the GP ranging from the glycan cap to the base (**Figure 6B, Figure S3**).

We then attempted to determine the critical binding residues at the amino acid level for a representative antibody from each of the 15 groups. Antibodies were screened for binding to alanine scanning mutant libraries of the EBOV GP. Screening was successful for EBOV-852, and we were able to identify single binding site residues for EBOV-598, EBOV-786, EBOV-709, and EBOV-823 (**Figure 6C**). However, as all these antibodies are neutralizing antibodies and therefore bind very avidly to the EBOV GP, single residue alanine mutations failed to disrupt binding for the rest of the antibodies even after digestion and screening of the binding of the antibodies as Fabs.

Critical residues for EBOV-852 were P279, E303, S302, and K299, all residues which span the glycan cap. These results are consistent with the competition-binding ELISA results, in which EBOV-852 competed with the glycan cap antibody 13C6 (**Figure 5A**). E303 is a conserved residue for not only EBOV, BDBV, and SUDV but also for TAFV and RESTV. S302 and P279 are also conserved between EBOV and BDBV. However, in SUDV and RESTV the serine is replaced with glycine and in SUDV only, the proline is replaced with alanine. These findings likely explain why EBOV-852 binds only to EBOV and BDBV GPs.

Critical residues for EBOV-598 are R89 and G149. The G149 residue is conserved across EBOV, BDBV, SUDV, TAFV, REST, and R89 is conserved across EBOV, BDBV, SUDV, TAFV, REST, and MARV. Although this residue sits in the conserved region of the receptor binding domain (RBD), the residue after it, S90 is only conserved between EBOV, SUDV, and REST. In BDBV, TAFV, and MARV, this residue is substituted to an alanine. Therefore, this finding likely explains the cross-reactive neutralization of EBOV and SUDV but not BDBV by EBOV-598, which was unexpected as EBOV and BDBV GP are generally more similar in sequence identity than EBOV and SUDV GP. A single critical residue was identified for EBOV-786, EBOV-709, and EBOV-823 (**Figure 6C**). The one critical residue indicated for EBOV-786 was S46. This residue is conserved between EBOV and BDBV but not SUDV GP (in which the serine changes to a threonine). However, this residue is also conserved in TAFV. Lastly, the critical residue identified for EBOV-709 and EBOV-823 is W275. This residue is conserved across EBOV, BDBV, SUDV, TAFV, and REST, and sits in the glycan cap, also mirroring the results of the competition-binding ELISA data as these antibodies compete with mAb 13C6 (**Figure 5A**) and the results of the negative stain EM studies (**Figure 6B**). Therefore, it is likely that these findings explain the capacity of EBOV-709 and EBOV-852 to neutralize EBOV and BDBV but not SUDV.

### Surveying the level of publicness in identified public clonotypes

There are many methods for identifying public clonotypes, using multiple identity thresholds or junction matching techniques. Our approach used a paired heavy and light chain gene sequence set and we characterized the functional phenotypes of all public clonotype antibodies identified, allowing us to use a sequence identity threshold on the lower end of common practice. Using the same clustering scheme of binning on the heavy chain V and J gene as well as CDR3 length, we next clustered the public clonotypes identified at 70% and 80% similarity on the CDR3 nucleotide sequence and binned them at the back end by matching on the light chain V and J gene. Next, we identified the antibodies that fell out of each public clonotype cluster at each threshold of 60%, 70%, and 80% and investigated if antibodies that fell out at each threshold shared similar functional phenotypes that would differentiate them from the main group (**Table S2**). We did not detect a difference in the functional binning of antibodies when clustering at differing identity thresholds.

Our criteria for public clonotype identification requiring the same heavy and light chains could be considered conservative, as there are numerous examples of public clonotypes defined by a recurrent heavy chain that undergo promiscuous pairing with various light chains (Setliff et al., 2018; Tan et al., 2021). To determine the flexibility of sequences on the light chain, we tested if public clonotypes within the same group would express and function with light chains belonging to differing donors. Three public clonotypes were selected for which the heavy chain from one donor and the light chain from another donor were recombined to investigate if the reactivity of the public clonotype was preserved. EBOV-1182 uses the heavy chain from EBOV-826 and the light chain from 2.1.1D07 and maintains its ability to neutralize both EBOV and BDBV when the antibody chains were swapped. EBOV-1190 uses the heavy chain from EBOV-786 and the light chain from 5.6.1A02 and neutralized both EBOV and BDBV. Lastly EBOV-1187 which uses the heavy chain from EBOV-852 and the light chain from 56-3-7A, also neutralizes both EBOV and BDBV (**Figure S2**). Together, we are confident that our approach of using a threshold of 60% on the CDRH3 sequence in conjunction with binning on the CDRH3 length and both heavy and light chain V and J genes is successful in identifying public clonotypes when using paired sequence sets.

### Germline-encoded properties are retained in some public clonotypes

To investigate if the neutralizing activity of these public clonotypes was due to germline-encoded reactivity or the result of somatic mutations, we investigated the equivalent germline-encoded antibodies for each public clonotype. We aligned each heavy and light chain variable region sequences to its respective germline gene sequence and reverted residue that differed from the germline gene to the inferred germline residue. Each germline-revertant (GR) antibody then was tested to see if the GR version of the antibody shared similar properties to its mutated counterparts (**Table S1**). All GR antibodies were tested for binding to EBOV, BDBV, or SUDV GP. Additionally, they were tested for neutralization of VSV-EBOV, -BDBV, or -SUDV. Although most GR antibodies did not retain binding to either GP or neutralize either virus, three GR antibodies retained functional activity compared to their mutated counterparts (**Figure 5A**). EBOV-852-GR, encoded by *IGHV1-2/IGLV2-8* retained ability to bind and neutralize EBOV and BDBV. EBOV-709-GR, encoded by *IGHV1-69/IGLV1-44* retained ability to bind and neutralize EBOV but only partially to BDBV (**Figure 5C**). EBOV-857, encoded by *IGHV3- 13/IGKV3-20* retained its ability to bind and neutralize EBOV (**Figure 5A**). These findings indicate that germline genes in these public clonotypes encode antibodies with critical residues that not only mediate binding but also neutralization.

### EBOV public clonotypes protect *in vivo*

We then tested these public clonotypes and their level of protection *in vivo* in mice against EBOV (Mayinga strain). Antibodies were delivered at 5 mg/kg 1 day after inoculation with EBOV. Scores on protection from death, weight loss, and disease were measured for 28 days. Treatment with mAbs representing public clonotypes conferred protection against mortality. 100% of animals survived the infection after treatment with EBOV-598, EBOV-790, EBOV-852, EBOV-705, EBOV-709, EBOV-801, EBOV-817, or EBOV-831. 80% of animals survived after treatment with EBOV-823 or EBOV-563, and 40% – after treatment with EBOV-822 (**Figure 7, Figure S4**). Although EBOV-854 showed low levels of neutralization *in vitro* using VSV-EBOV, it did not show neutralization with authentic virus, and accordingly failed to protect animals *in vivo*. Overall, these findings show that there are public clonotypes specific to EBOV that protect *in vivo*.

**Figure 7.**
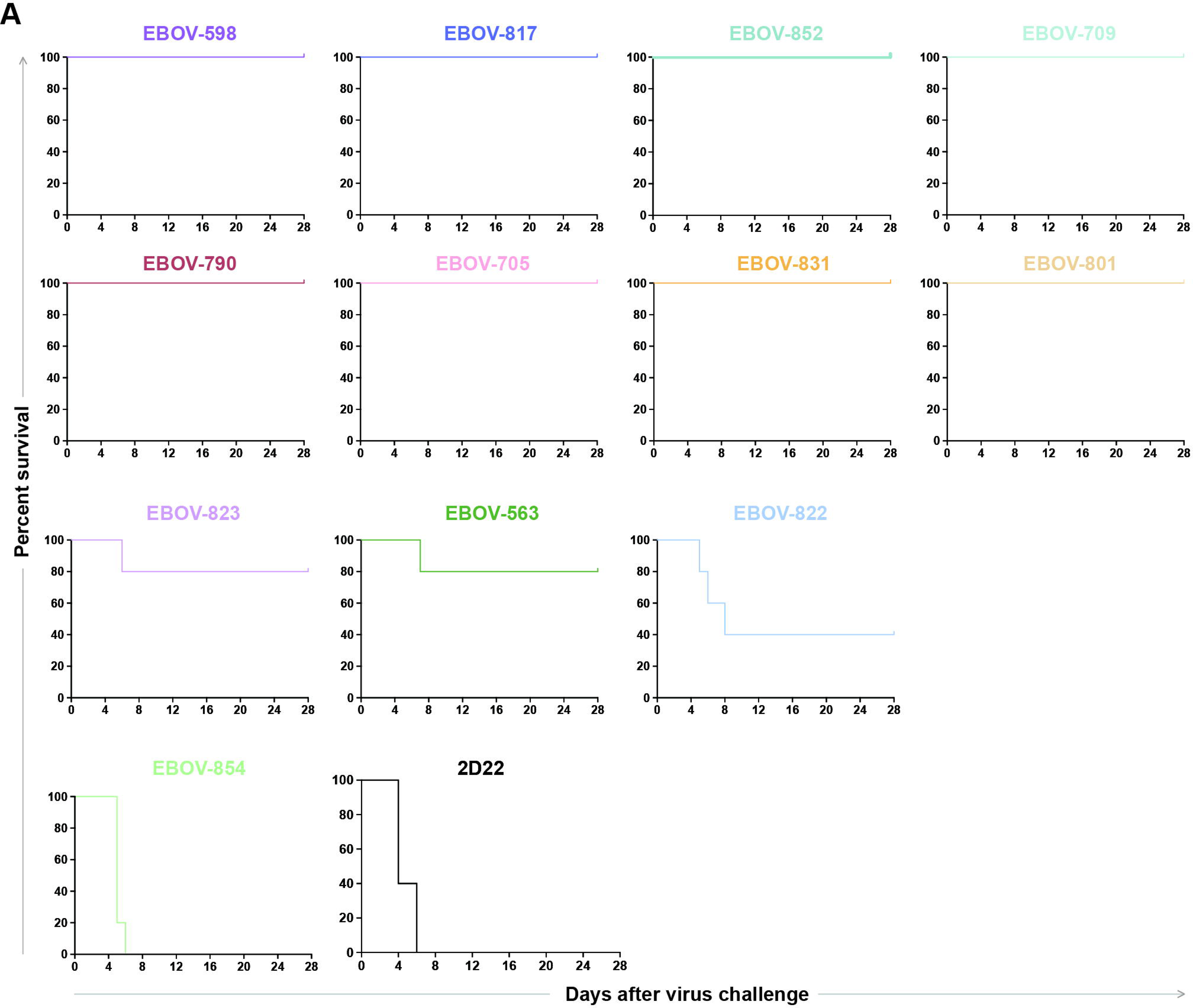
*In vivo* protection using public clonotypes. Mice (n = 5) were treated i.p. with 100 µg (∼5 mg/kg) of an individual antibody per mouse on day 1 post-challenge. Human antibody DENV 2D22 (specific to dengue virus) served as a negative control. Mice were monitored twice daily from day 0 to day 14 post challenge for survival and monitored daily from day 15 to 28 as described previously (Ilinykh et al., 2018).

## DISCUSSION

The size of the human B cell response to a pathogenic virus protective antigen has not been defined. Previous work has established that the overall circulating repertoire of each individual contains around 11 million or more B cell clonotypes defined by V_H_, J_H_, and CDR3 amino acid sequence using bulk sequencing data (Briney et al., 2019; Soto et al., 2019), however, the number and diversity of B cells specific to viral antigens is poorly understood at the paired sequence level. Here we present the largest individual antigen-specific repertoire from a single sample reported, to estimate the size and complexity of an individual’s response to a virus. After sorting 100,000 GP-specific B cells, we recovered paired antibody genes for over 15,000 clones and found over 10,000 clonotypes in that repertoire. Species richness calculations estimate that the individual’s sample contained over 20,000 clonotypes reactive with EBOV GP at the time point tested, about 9 months after infection. It should be noted that each of those >20,000 clonotypes contains many somatic variants (for instance, the largest clonal family we recovered had 21 sequences within its single lineage in this study). Thus, the size and complexity of the response to a single viral protein is enormous.

A strength of the antibody discovery approach used here was that we not only obtained variable gene sequences, but also those sequences were authentically paired heavy and light chain sequences from single cells. This approach allowed us to express representative naturally occurring mAbs of each of the clonotypes of interest so that we could validate their specificity and define their cross-reactivity and neutralizing potency. Here, we observed that 45% of the clonally expanded antibody repertoire neutralized EBOV. About two-thirds of the neutralizing clones targeted the glycan cap region of the GP. This finding shows that, even though many have considered the glycan cap a poor target for protective responses, most neutralizing antibodies in the clonally expanded repertoire target the glycan cap. Additionally, we mapped functional characteristics of each clonally expanded family to genetic similarities of antibodies not only within clonal families but also between clonal families at a scale previously unseen, allowing for visualization of clustering of functionally similar antibodies with genetically similar CDR3s.

The nature of future ebolavirus epidemics cannot be predicted, and therefore it is important to understand how cross-reactive neutralizing antibodies arise in response to the virus. For the cross-reactive neutralizing antibodies identified that recognized multiple ebolavirus species, we investigated if that cross-reactive neutralizing activity was germline-encoded or acquired through acquisition of somatic mutations. The studies of UCAs revealed that the *IGHV1-69* and *IGHV1-2* heavy chain variable gene segments can encode cross-reactive neutralizing antibodies, suggesting the origin of heterologous immunity in individuals infected with one ebolavirus species.

Public clonotypes have been identified in human antibody repertoires in response to a variety of viral pathogens including influenza virus (Joyce et al., 2016; Pappas et al., 2014; Zost et al., 2021), respiratory syncytial virus (Mukhamedova et al., 2021), hepatitis C virus (Bailey et al., 2017), HIV (Setliff et al., 2018; Zhou et al., 2015), and SARS-CoV-2 (Chen et al., 2021; Sakharkar et al., 2021; Schmitz et al., 2021; Tan et al., 2021) revealing selection of genetically similar B cell receptors in memory cells in circulation of diverse immune individuals. Understanding the prevalence of public clonotypes and their functionalities requires very large numbers of paired sequences.

Several public clonotypes have previously been reported that recognize the EBOV GP (Cagigi et al., 2018; Cohen-Dvashi et al., 2020; Davis et al., 2019; Rijal et al., 2019; Wec et al., 2017). But the large scale of sequencing obtained in this study uniquely positioned us to systematically identify a high prevalence of public clonotypes elicited to the EBOV GP, with 73 public clonotypes, a level of sharing that is unexpected since we required the public clonotypes to share not only heavy chain features but also the same light chain gene usage. This data collection is the largest set of B cell public clonotypes reported to date for a viral pathogen, and most of these are novel public clonotypes to EBOV that have not yet been described. By functionally characterizing every antibody identified in the 73 public clonotypes, we found that roughly 20% of EBOV GP specific public clonotypes neutralized the virus. Most of the neutralizing public clonotypes also conferred therapeutic protection *in vivo* against lethal challenge. The studies using negative stain electron microscopy revealed that these 15 neutralizing public clonotypes target diverse regions of the GP ranging from the glycan cap to the base. Additionally, analysis into the germline-encoded functions of public clonotypes revealed that three of the 15 neutralizing public clonotypes retained neutralization when somatic mutations were reverted to the inferred germline gene segment sequence. Public clonotypes that neutralized virus as UCA antibodies were encoded by *IGHV1-69*, *IGHV1-02*, or *IGHV3-13*.

Mining for and understanding the properties of public clonotypes informs a deeper understanding of population immunity by revealing immunodominant B cell responses within immune populations, which may be of benefit for rational design of vaccines that may exhibit immunogenicity in a broader segment of the population. Knowing the public clonotype profile following natural infection also can enhance experimental vaccine testing, since the immunogenicity for desirable public antibodies recognizing cross-reactive sites of vulnerability for potent neutralization can be recognized at the cDNA sequence level. We should also keep in mind, however, that the broad induction of public antibody clonotypes recognizing the protective antigen of an RNA virus can lead to a constant and collective pressure on certain epitopes to viruses leading to rapid selection of escape mutant variants.

The large size of the data set of EBOV GP-reactive memory B cells created in this study provided an opportunity to mine for public clonotypes specific to EBOV GP but also posed technical challenges for data analysis. We sought to identify public clonotypes using both heavy and light chain sequences, a workflow that is only achievable with paired sequences from single B cells. Identity thresholds used to identify public clonotypes in variable gene sequence sets vary greatly in the field. We chose an identity threshold for identification of public clonotypes on the lower end of common conventions, but our confidence in these assignments was supported by both heavy and light chain gene segment assignments and with functional testing of recombinant antibodies encoded by these sequences. As we tested all 294 members of the public clonotypes functionally, we were in a unique position to investigate different clustering thresholds for identifying public clonotypes and tested how those thresholds affected the grouping of antibodies with functional phenotypes. As clustering at higher thresholds did not necessarily bin antibodies into tidier functional phenotype bins, we conclude that when mining for public clonotypes within an antigen-specific sequence set using paired sequencing from single cells, a threshold of 60% identity in the CDRH3 is sufficient.

It is highly desirable to identity germline-encoded pan-ebolavirus cross-reactive antibodies to support rational vaccine design and testing efforts. Here we identified two V_H_ genes that often encode cross-reactive antibodies at the germline level: *IGHV1-69* and *IGHV1-02*. For *IGHV1-69*, a structural explanation is now possible. *IGHV1-69/IGHJ6-*encoded antibodies have been described to target the mucin-like domain (MLD) cradle, exploiting hydrophobic residues encoded by the germline of these gene segments bind and destabilizes the GP quaternary structure and therefore blocking cleavage required for receptor binding (Murin et al., 2021). Within the antibodies described, potency was acquired through somatic hypermutation. However, antibodies in this class had cross-reactive neutralizing proprieties regardless since they target the conserved MLD anchor and cradle. The molecular and structural determinants of the cross-reactive activities associated with *IGHV1-02-*encoded antibodies have yet to be described. Here, the data reveal that the cross-reactive neutralizing properties of antibodies encoded by *IGHV1-02* are germline-encoded. *IGHV1-02* also has been shown to encode broadly neutralizing antibodies for HIV, due to hydrophobic residues in the CDR2 similar to those encoded by the CDR2 of some alleles of the germline gene *IGHV1-69*. As all the *IGHV1-02*-encoded antibodies discovered here competed for binding with the glycan cap mAb 13C6, we predict that *IGHV1-02* encodes for cross-reactive neutralizing antibodies targeting the glycan cap or MLD region of the GP using a similar mechanism to that of *IGHV1-69*-encoded antibodies.

Antibodies encoded by *IGHV1-69* and *IGHV1-02* may represent a substantial portion of first-line of defense during ebolavirus infection. We speculate that the earliest neutralizing response to ebolavirus infection is likely encoded *IGHV1-69* and *IGHV1-02*, since all antibodies discovered here for which the UCA antibodies neutralized virus competed for binding with the glycan cap mAb 13C6. When identifying the breakdown of serum IgG protein antibodies, 10% of those identified used *IGHV1-02* and 7% used *IGHV1-69*. Additionally, these serum antibodies mapped back to large clonally expanded families that included clonal families with 13 or 20 members respectively.

It has been reported that B cells circulating early in convalescence target the glycan cap region of the GP (Williamson et al., 2019). We found that 9.2% of EBOV GP-specific antibodies in this large repertoire were encoded by *IGHV1-69*, causing it to be the most heavily used variable gene, with the *IGHV4-34* or *IGHV1-02* genes also used frequently, at 7.5% or 6.3%, respectively. This finding further demonstrates a substantial reliance on germline-encoded antibody responses in the humoral immune response to EBOV. The intrinsic hydrophobic properties of antibodies encoded by these genes likely play a vital role in immunity to ebolaviruses and other viruses.

Antibodies are not secreted by circulating memory B cells but rather by long-lived plasma cells in the bone marrow. We have recently described the antibody response in convalescent plasma within the same donor (Gilchuk et al., 2021). However, little is known about the diversity of antibodies that overlap between the memory B cell repertoire and the plasma antibody repertoire. Previous data had shown that in plasma, there is preferential recognition of the cleaved EBOV GP. However, in our study, it is interesting that when narrowing the antibody characteristics of plasma antibodies identified from clonally expanded families, these antibodies preferentially recognized the intact GP. Therefore, it seems likely that the bulk of plasma antibodies could derive from specificities less common in circulating memory B cells (noted as singlets in our single cell RNAseq repertoire) where a lot of the reactivity for cleaved GP would reside. A technical limitation of plasma proteomic antibody studies is that likely only highly represented antibodies are detected, and therefore this dataset is likely lacking antibodies present at low levels in the plasma.

This in-depth systematic study in both the private and public antibody response to EBOV GP uniquely ties the importance of repertoire wide studies with functional studies of mAbs. Understanding which germline gene combinations may encode neutralizing antibodies and detailing the relationship between antibody gene usage and the corresponding immunoglobulin antiviral breadth and function also could be used in future machine learning and artificial intelligence studies for *in silico* discovery of immunogenic vaccines and new broad and potent antibodies. This general approach also could be generalizable to understanding the humoral response to other pathogenic viruses at a repertoire-wide level.

## Supplemental information

Supplemental information including supplemental figures and tables can be found with this article online.

## Acknowledgments

EM data collections were conducted at the Center for Structural Biology Cryo-EM Facility at Vanderbilt University. We thank Dr. Cinque Soto for advice and review of the bioinformatic approaches used in the paper and Rachael Nargi for excellent technical support.

## Author contributions

Conceptualization, E.C.C. and J.E.C.; Investigation, E.C.C, P.G., S.J.Z., E.B., P.A.I., K.H., L.M., S.B., S.D., C.K., A.T., J.R., R.S., R.N., E.D., E.O.S.; Writing first draft: E.C.C. and J.E.C; All authors edited the manuscript and approved the final submission; Supervision, B.J.D., N.C., A.B., R.H.C., J.E.C.; Funding acquisition, A.B., J.E.C.

## Declaration of interests

E.D. and B.J.D. are employees of Integral Molecular, and B.J.D.is a shareholder in that company. J.E.C. has served as a consultant for Eli Lilly, GlaxoSmithKline and Luna Biologics, is a member of the Scientific Advisory Board of Meissa Vaccines and is Founder of IDBiologics. The Crowe laboratory has received funding support in sponsored research agreements from AstraZeneca, IDBiologics, and Takeda. All other authors declare no competing interests.

**Figure S1. Functional characteristics of the EBOV GP-specific clonally-expanded repertoire**. The functional characteristics of all 224 clonally-expanded antibodies are listed in this table. The first column shows the number of antibodies in the cluster, the second column shows the CDRH3 amino acid length, the third column shows the CDRL3 amino acid length. Blue boxes indicate binding to GPs in ELISA, purple indicates binding in a cell-surface GP display assay. Green indicates neutralizing activity for VSV-EBOV, and pink indicates lack of detectable neutralizing activity.

**Table S2. Sequences of the unmutated common ancestors and germline revertant antibodies.** Sequences of unmutated common ancestors for the clonally-expanded antibodies and the germline revertant antibodies from the public clonotypes are listed.

**Figure S3. Functionality of public clonotypes after swapping heavy and light chains.** Heavy chains from a public clonotype antibody from one donor and light chain from that public clonotype observed in another donor were paired and expressed together and tested for neutralization of VSV-EBOV, -BDBV, or -SUDV.

**Figure S4. Negative stain electron microscopy complexes of representative antibodies from each public clonotype.** IgG for a representative antibody for each public clonotype was digested to obtain the Fab form of antibody and complexed with EBOV GP.

**Figure S5. *In vivo* efficacy of public clonotypes.** Mice (n = 5) were treated i.p. with 100 µg (∼5 mg/kg) of individual antibody per mouse on day 1 post-challenge. MAb DENV 2D22 was used as a negative control. Mice were monitored twice daily from day 0 to day 14 post-challenge for illness, survival, and weight loss, followed by once daily monitoring from day 15 to the end of the study at day 28.

a. Illness scores of mice treated with each public clonotype.
b. Body weight graphs of mice treated with each public clonotype.

**Table S1. Sequences of the unmutated common ancestors and germline revertant antibodies.** Sequences of unmutated common ancestors for the clonally-expanded antibodies and the germline revertant antibodies from the public clonotypes are listed.

**Table S2. Functional characteristics of all public clonotypes.** Functional testing for each of the 294 public clonotypes tested are listed. Clustering threshold analysis is also included in this table.

## STAR METHODS

### Research participants

Human PBMCs and plasma were obtained at Vanderbilt University Medical Center in Nashville, TN, USA, from a survivor of the 2014 EVD epidemic after written informed consent. The studies were approved by the Vanderbilt University Medical Center Institutional Review Board. PBMCs and plasma were collected after the illness had resolved. The donor is a male human survivor of the 2014 EVD outbreak in Nigeria and was 31 years of age when infected, and 32 when PBMCs and plasma were collected 15 months later. At the time of blood collection, plasma samples were tested by qRT-PCR and found to be negative for the presence of viral RNA.

### Cell lines

Vero-E6, Jurkat, Vero CCL-81, and THP-1 cells were obtained from the American Type Culture Collection (ATCC). Vero-E6 cells were cultured in Minimal Essential Medium (MEM) (Thermo Fisher Scientific) supplemented with 10% fetal bovine serum (FBS; HyClone) and 1% penicillin-streptomycin in 5% CO_2_, at 37°C. ExpiCHO (hamster, female origin) and FreeStyle 293F cell lines were purchased from Thermo Fisher Scientific and cultured according to the manufacturer’s protocol. The Jurkat-EBOV GP (Makona variant) cell line stably transduced to display EBOV GP on the surface(Davis et al., 2019) was a kind gift from Carl Davis (Emory University, Atlanta, GA). Jurkat-EBOV GP and THP-1 cells were cultured in RPMI 1640 (Gibco) medium supplemented with 10% FBS and 1% penicillin-streptomycin in 5% CO2, at 37°C.

### Viruses

The mouse-adapted EBOV Mayinga variant (EBOV-MA, GenBank: AF49101) (Bray et al., 1998), authentic EBOV Mayinga variant expressing eGFP(Towner et al., 2005), infectious vesicular stomatitis virus rVSV/EBOV GP (Mayinga variant), rVSV/BDBV GP, rVSV/SUDV GP(Garbutt et al., 2004), and chimeric EBOV/BDBV-GP and EBOV/SUDV-GP(Ilinykh et al., 2016) were used for mouse challenge studies or neutralization assays. Viruses were grown and titrated in Vero-E6 cell monolayer cultures.

### GP expression and purification

For B cell labeling, and flow cytometric sorting, we used EBOV GP produced in Drosophila Schneider 2 (S2) cells. Briefly recombinant ectodomain of EBOV GP Δ in a modified pMTpuro vector was transfected into S2 cells followed by stable selection of transfected cells with 6 µg/mL of puromycin. GP ectodomain expression was induced with 0.5mM CuSO_4_ for 4 days. Protein was purified using Step-Tactin resin (Qiagen) via an engineered strep II tag and purified further by Superdex 200 (S200) column chromatography. For ELISA studies, the ectodomains of EBOV GP ΔTM (residues 1-636; strain Makona; GenBank KM233070), BDBV GP ΔTM (residues 1- 643; strain 200706291 Uganda; GenBank: NC_014373), SUDV GP ΔTM (residues 1-637; strain Gulu; GenBank: NC_006432), and MARV GP ΔTM (residues 1-648; strain Angola2005; GenBank: DQ447653) were expressed using the FreeStyle 293F cell line and purified as described (Gilchuk et al., 2018).

### Memory B cell isolation and flow cytometric analysis

PBMCs from a leukopak were isolated with Ficoll-Histopaque by density gradient centrifugation. The cells were cryopreserved in the vapor phase of liquid nitrogen until use. Total B cells were enriched by negative selection from PBMCs using EasySep Human Pan-B Cell Enrichment Kit (StemCell Technologies). Enriched cells were stained on ice in Robosep buffer containing the following phenotyping antibodies: anti-Human CD19-PE, anti-IgM-FITC, anti-IgD-FITC. The EBOV GP-reactive memory B cells were labeled with recombinant EBOV GP protein that was produced in Drosophila S2 cells as described above and purified by flow cytometric cell sorting using an SH800 cell sorter (Sony) as described previously (Gilchuk et al., 2020b). Approximately 100,000 cells were FACS-sorted in bulk for downstream paired antibody heavy and light chain variable gene sequence analysis.

### Generation of antibody variable-gene libraries from single B cells

For paired antibody variable gene sequence analysis, cells were resuspended into DPBS containing 0.04% non-acetylated BSA, split into four replicates, and separately added to 50 μL of RT Reagent Mix, 5.9 μL of Poly-dt RT Primer, 2.4 μL of Additive A and 10 μL of RT Enzyme Mix B to complete the Reaction Mix as per the vendor’s protocol. The reactions then were loaded onto a Chromium chip (10x Genomics). Chromium Single Cell V(D)J B-Cell-enriched libraries were generated, quantified, normalized and sequenced according to the User Guide for Chromium Single Cell V(D)J Reagents kit (CG000086_REV C). Amplicons were sequenced on an Illumina Novaseq 6000, and data were processed using the CellRanger software v3.1.0 (10x Genomics).

### Bulk sequence analysis of antibody variable region genes

Total RNA was extracted from approximately 5,000 B cells using the Qiagen RNeasy Micro kit following the manufacturer’s recommendations (Qiagen). To maximize target enrichment recovery, we employed two separate library preparation approaches with three separate primer mixes to avoid any individual primer set’s amplification bias. In the first library preparation approach, we used the OneStep SuperScript III Platinum^®^Taq High Fidelity kit (Thermo Fisher Scientific) in a one-step RT-PCR approach with 2 µL total RNA as input into separate reactions to enrich for B cell heavy- and light-chain transcripts. In the first set of one-step RT-PCR reactions we used a combination of previously published primer sets (Diss et al., 2002; Smith et al., 2009; van Dongen et al., 2003), while in the second set of reactions we used in-house designed heavy- and light-chain primers targeting the beginning or end of FR1 or FR4 of B cell transcripts, respectively. All primer sequences used for the one-step RT-PCR approach are listed below. The thermal cycling parameters for both sets of reactions were as follows: 50°C for 30 min; 94°C for 2 min; 24 cycles of 94°C for 15 s, 58°C for 30 s, and 68°C for 1 min; 68°C for 10 min. PCR products were purified using a 2% gel cassette on a PippinHT system (Safe Science) targeting 200-500 bp amplicons (Sage Science). Illumina indexing and adapter ligation was performed using the NEBNext® Ultra DNA Library Prep kit (NEB).

In the second library preparation approach, 4 µL total RNA was shipped to and processed by AbHelix, LLC (www.abhelix.com, South Plainfield, NJ, USA). Briefly, RNA samples were reverse-transcribed using oligo d(T) 18 and SuperScript IV Reverse Transcriptase (Thermo Fisher Scientific) followed by Ampure XP bead purification (Beckman Coulter). The purified RT products were divided evenly for the first round of PCR amplification specific to human IgG, IgK, IgL, IgM, or IgA. The 5′ multiplex PCR primers are designed within the leader sequences of each productive V-gene and the 3′ primers within the constant regions but in close approximation to the J-C junctions. The resulting first-round PCR products were purified with magnetic beads and subjected to the second round of PCR amplification to add Illumina index and adapter sequences. The resulting PCR products were purified with Ampure XP (Beckman Coulter) magnetic beads and pooled. Phusion High-Fidelity DNA Polymerase (Thermo Fisher Scientific, CA) was used in all PCR amplification reactions and care was taken to minimize the number of cycles to achieve adequate amplification. Primer sequences used by Abhelix are proprietary and are not provided here.

The DNAs in the final resulting libraries from both library preparation approaches were quantified using the Qubit 3.0 fluorometer (Thermo Fisher Scientific) prior to size determination using a Bioanalyzer 2100 (Agilent). Libraries were re-quantified using the KAPA qPCR kit (KAPA Biosystems) before sequencing on an MiSeq instrument (Illumina) using two separate 2 x 300 bp flow cells (Illumina).

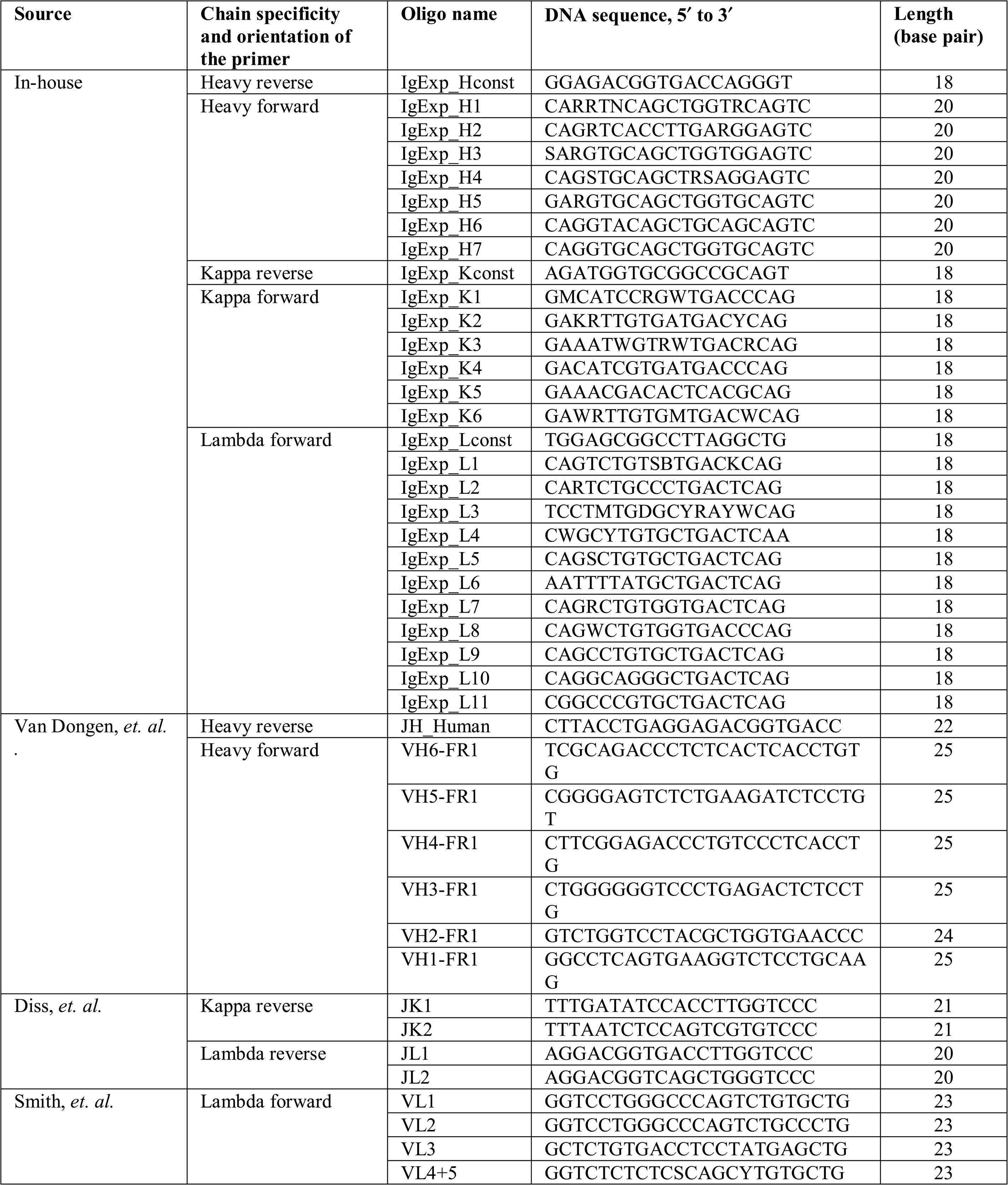
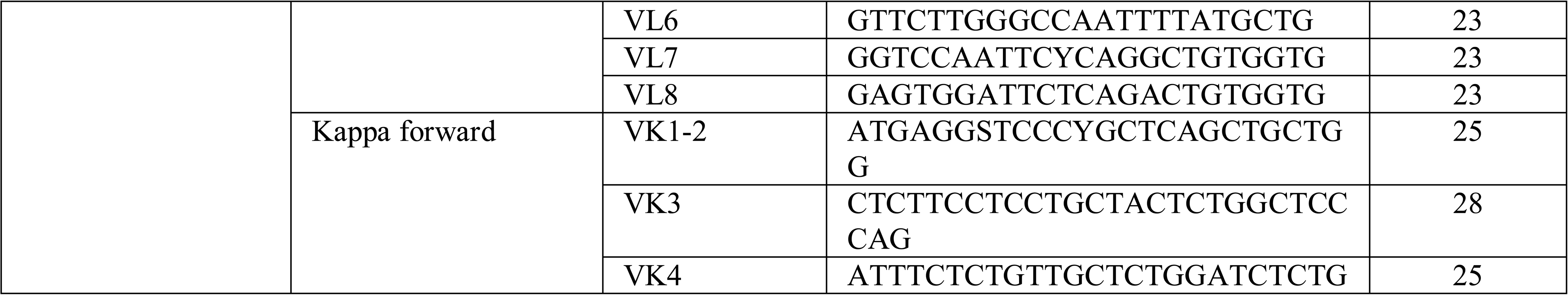

### Paired sequence clustering

To identify clonal families, paired sequences obtained from our antigen specific sort was obtained. Sequences were then clustered based on genetic similarity. Sequences were first binned together if they shared the same heavy chain V and J gene as well as CDRH3 length. After, sequences were clustered according to 80% sequence similarity on the CDRH3 nucleotide sequence. Then, they were binned together if they shared the same light chain V and J gene as well as CDRL3 length. Lastly, sequences were clustered again according to 80% sequence similarity on the CDRL3 sequence. These resulting clusters of sequences were designated as clonal families.

To identify public clonotypes, publicly available paired sequence sets of antibody genes were obtained (Bornholdt et al., 2016; Davis et al., 2019; Ehrhardt et al., 2019; Rijal et al., 2019). Together with sequences derived from this paper, public clonotypes were determined by genetic similarities of antibody sequences using the following clustering scheme. They were first binned by V_H_ and J_H_ gene and CDRH3 amino acid length. Sequences within each bin then were clustered according to 60% sequence similarity on their CDRH3 nucleotide sequence. Lastly, sequences were binned if they used the same light chain V and J gene. Clusters of sequences meeting the described criteria and contained sequences originating from two or more individuals were deemed public clonotypes.

### Bulk sequence clustering

Sequences within a same paired sequence cluster was taken. These sequences were then used to search for sequences within the bulk sequence dataset. Sequences sharing the same V and J gene as well as 80% similarity on the CDR3 sequence were then clustered together.

### Heat map generation

The heavy chain variable gene and light chain variable gene used for each public clonotype were tallied. The number of public clonotypes with corresponding V_H_-V_L_ genes were counted. These frequency counts were then plotted onto the heatmap using Python Seaborn Library.

### Network generation

Antibody sequences within the same clonal family was taken to compute a centroid sequence using Vsearch v2.7.1 to be used as a representative for that clonal family. The hamming distance of each antibody sequence within the clonal family to its respective centroid was then calculated. The distance between centroids belonging to different clonal families were then calculated using Levenshtein distance. Distances were calculated using the Python distance library (https://pypi.org/project/Distance/) for hamming distance. Levenshtein distance was calculated as described in literature(Miho et al., 2019). The graph was created with NetworkX and visualized using Matplotlib and PyGraphviz.

### Species richness calculations

Clonal families identified as described above, were utilized as a taxonomic unit/species. Rarefaction curves were calculated based on clonal families and unique members as species and individuals respectively for 10,000 repetitions with RTK (Saary et al., 2017). The mean values of these repetitions were plotted for species richness and the Chao1 estimate of abundance. Fluctuations and rise in Chao1 estimate for the non-antigen specific data set are interpreted to mean that sequencing depth was inadequate to capture an accurate estimate.

### Construction of maximum likelihood trees

Sequences belonging to each cluster/clonal family were aligned to their corresponding germline gene using Clustal Omega v1.2.0. We used the PHYLIP phylogenetic software package v3.697 to generate maximum-likelihood trees from the aligned sequences using the DNAML program, using the sequence of the germline IGHV or IGKV/IGLV as an out group. The resulting phylogenetic trees were visualized using Geneious Prime v2019.2.1. Branches were colored corresponding to the sequence set in which they were identified. The inferred unmutated common ancestor (UCA) was extracted from the PHYLIP-generated tree.

### Antibody production and purification

Sequences of mAbs were synthesized using a rapid high-throughput cDNA synthesis platform (Twist Bioscience) and subsequently cloned into an IgG1 monocistronic expression vector (designated as pTwist-mCis_G1) for mAb secretion from mammalian cell culture. This vector contains an enhanced 2A sequence and GSG linker that allows simultaneous expression of mAb heavy- and light-chain genes from a single construct upon transfection (Chng et al., 2015). We performed transfections of ExpiCHO cell cultures using the Gibco ExpiCHO Expression System and protocol for 50 mL mini bioreactor tubes (Corning) as described by the vendor. Culture supernatants were purified using HiTrap MabSelect SuRe (Cytiva) on a 24-column parallel protein chromatography system (Protein Biosolutions). Purified mAbs were buffer-exchanged into PBS, concentrated using Amicon Ultra-4 50-kDa centrifugal filter units (Millipore Sigma) and stored at 4°C until use.

### ELISA binding assays

Wells of 384-well microtiter plates were coated with purified recombinant GP at 4°C overnight. Plates were blocked with 2% non-fat dry milk and 2% normal goat serum in DPBS containing 0.05% Tween-20 for 1 h. All antibodies were diluted to a concentration of either 0.4 µg/mL for the matured antibodies or 5 µg/mL for the germline-revertant antibodies. Antibodies were diluted in two-fold dilutions until binding was no longer detected. Bound antibodies were detected using goat anti-human IgG conjugated with horseradish peroxidase and TMB substrate. The reaction was quenched with 1N hydrochloric acid once color was developed. The absorbance was measured at 450 nm using a spectrophotometer (Biotek).

### Real-time cell analysis (RTCA) neutralization assay

To determine neutralizing activity of purified antibodies or human serum, we used real-time cell analysis (RTCA) assay on an xCELLigence RTCA MP Analyzer (ACEA Biosciences Inc.) that measures virus-induced cytopathic effect (CPE) (Gilchuk et al., 2020a; Suryadevara et al., 2021; Zost et al., 2020). Briefly, 50 μL of cell culture medium (DMEM supplemented with 2% FBS) was added to each well of a 96-well E-plate to obtain background reading. A suspension of 15,000 Vero cells in 50 μL of cell culture medium was seeded in each well, and the plate was placed on the analyzer. Measurements were taken automatically every 15 min, and the sensograms were visualized using RTCA software version 2.1.0 (ACEA Biosciences Inc). VSV-EBOV, VSV-BDBV, and VSV-SUDV were mixed 1:1 with a respective dilution of mAb using DMEM supplemented with 2% FBS as a diluent and incubated for 1 h at 37°C in 5% CO_2_. At 16 h after seeding the cells, the virus-mAb mixtures were added in replicates to the cells in 96-well E-plates. Triplicate wells containing virus only (maximal CPE in the absence of mAb) and wells containing only Vero cells in medium (no-CPE wells) were included as controls. Plates were measured continuously (every 15 min) for 48 h to assess virus neutralization. Normalized cellular index (CI) values at the endpoint (48 h after incubation with the virus) were determined using the RTCA software version 2.1.0 (ACEA Biosciences Inc.). Results are expressed as percent neutralization (CI of wells divided by CI of cells only wells) in a presence of respective mAb relative to control wells with no CPE minus CI values from control wells with maximum CPE. RTCA IC_50_ values were determined by nonlinear regression analysis using GraphPad Prism 9 software.

### Competition-binding ELISA

Wells of 384-well microtiter plates were coated with purified recombinant EBOV GP at 4°C overnight. Plates were blocked with 2% bovine serum albumin (BSA) in DPBS containing 0.05% Tween-20 for 1 h. Each antibody was diluted to a concentration of 10 µg/mL. Next, biotinylated antibodies were diluted to 2.5 µg/mL and added to the primary antibody solution without washing the plate to a final concentration of 0.5 µg/mL. Biotinylated antibody binding was detected with horseradish peroxidase-conjugated avidin (Sigma) and developed with TMB. The reaction was quenched with 1N hydrochloric acid once color was developed. Absorbance was measured at 450 nm using a spectrophotometer.

### Cell-surface binding to cleaved or intact GP

Alexa Fluor 647 NHS ester (Thermo Fisher Scientific) was used for antibody labeling. Binding of purified polyclonal or monoclonal antibodies to Jurkat-EBOV GP or Jurkat-EBOV GPCL cells was assessed by flow cytometry using an iQue Screener Plus high throughput flow cytometer (Intellicyt Corp.) as described previously (Gilchuk et al., 2018; Gilchuk et al., 2020b). Briefly, 50,000 cells were added per each well of V-bottom 96-well plate (Corning) in 5 mL of the DPBS containing 2% heat-inactivated ultra-low IgG FBS (Gibco) (designated as incubation buffer). Serial dilutions of antibody were added to the cells in replicates for a total volume of 50 µL per well, followed by 1 h incubation at ambient temperature, or 4°C in some experiments. Unbound antibody was removed by washing with 200 µL of the incubation buffer. Staining of cells was measured by flow cytometric analysis using the IntelliCyt iQue Screener Plus. Data for up to 20,000 events were acquired, and data were analyzed with ForeCyt (Intellicyt Corp.) software. Dead cells were excluded from the analysis based on forward and side scatter gates to identify the viable cell population. Binding to un-transduced Jurkat cells or binding of dengue antigen-specific mAb DENV 2D22 served as negative controls for most experiments.

Cells that displayed cleaved GP were prepared as described previously (Davis et al., 2019; Gilchuk et al., 2018; Gilchuk et al., 2020b). Briefly, Jurkat-EBOV GP cells were washed with DPBS containing calcium and magnesium (DPBS++), resuspended at 10^6^ cells/mL in DPBS containing 0.5 mg/mL of thermolysin (Promega), and incubated for 20 min at 37°C. The cleavage reaction was inhibited by washing cells with the incubation buffer containing DPBS, 2% of heat-inactivated FBS and 2 mM EDTA (pH 8.0). The GP cleavage was confirmed by loss of mAb 13C6 binding and high-level of binding that assessed with RBD-specific mAb MR78 relative to intact Jurkat-EBOV GP antibody binding. Antibody binding to un-transduced Jurkat (mock) cells served as a control for specificity of antibody staining. For screening of the micro-scale purified mAbs, cells were incubated with individual mAbs at a single 1:10 dilution, and the bound antibodies were detected using goat anti-human IgG antibody conjugated with PE (Southern Biotech).

### Mouse challenge with EBOV

Mice were housed in microisolator cages and provided food and water *ad libitum*. Groups of 7-8-week-old BALB/c mice (Charles River Laboratories) were inoculated with 1,000 plaque-forming units of EBOV-MA by the intraperitoneal (i.p.) route. Mice (n = 5) were treated i.p. with 100 μg (∼5 mg/kg) of individual mAb per mouse on day 1 post-challenge. Human mAb DENV 2D22 (specific to dengue virus) served as negative control. Mice were monitored twice daily from day 0 to day 14 post-challenge for illness, survival, and weight loss, followed by once daily monitoring from day 15 to the end of the study at day 28, as described elsewhere (Ilinykh et al., 2018). Moribund mice were euthanized as per the IACUC-approved protocol. All mice were euthanized on day 28 after EBOV challenge.

### Neutralization assay

Neutralization was tested against GFP-expressing EBOV and chimeric EBOV/BDBV-GP and EBOV/SUDV-GP constructs in a high-throughput format, as previously described (Ilinykh et al., 2016). The neutralization assays were performed using Vero-E6 cells. Neutralization assays were performed in triplicate, across 12 four-fold dilutions, starting from 200 μg/mL.

### Electron microscopy sample and grid preparation, imaging and processing of EBOV GP– Fab complexes

For electron microscopy imaging of EBOV GP and Fabs, Fabs were produced by digesting recombinant chromatography-purified IgGs using resin-immobilized cysteine protease enzyme (FabALACTICA, Genovis). The digestion occurred in 100 mM sodium phosphate and 150 mM NaCl pH 7.2 (PBS) for around 16 h at ambient temperature. To remove cleaved Fc from intact IgG, the digestion mix was incubated with CaptureSelect Fc resin (Genovis) for 30 min at ambient temperature in PBS buffer. For screening and imaging of negatively-stained EBOV protein in complex with human Fabs, the proteins were incubated at a Fab:EBOV GP (trimer) molar ratio of 4:1 for about 1 hour at ambient temperature, and approximately 3 μL of the sample at concentrations of about 10 to 15 μ/mL was applied to a glow-discharged grid with continuous carbon film on 400 square mesh copper electron microscopy grids (Electron Microscopy Sciences). The grids were stained with 0.75% uranyl formate (Ohi et al., 2004) . Images were recorded on a Gatan US4000 4k×4k CCD camera using an FEI TF20 (TFS) transmission electron microscope operated at 200 keV and control with SerialEM (Mastronarde, 2005) All images were taken at 50,000× magnification with a pixel size of 2.18 Å per pixel in low-dose mode at a defocus of 1.5 to 1.8 µm. The total dose for the micrographs was around 30 e−/per Å^2^. Image processing was performed using the cryoSPARC software package (Punjani et al., 2017) . Images were imported, CTF-estimated with CTFFIND4 (Rohou and Grigorieff, 2015) and particles were picked automatically with template picker (a part of cryoSPARC). The particles were extracted with a box size of 160 pixels and binned to 80 pixels (pixel size of 4.36 Å/pix). Multiple 2D class averages were performed, and good classes were selected for *ab-initio* 3D map reconstruction. At the final step, the data sets were refined. Maps were imaged using Chimera software (Pettersen et al., 2004).

### Proteogenomic analysis

The immunoproteogenomic platform Alicanto (Bonissone, 2021) was used for identifying antibody sequences and visualizing proteomics results, similar to a previous study(Gilchuk et al., 2021). Briefly, the variable region sequences of antigen-sorted and sequenced B cells were analyzed and annotated by Alicanto. The tandem mass spectra were searched against this custom antibody database. Antibody clones were determined as present if unique peptide coverage exceeded 50% of the CDR3 region and general peptide coverage was 100%, while coverage over the entire variable region sequence was above 90%.

### Quantification and statistical analysis

The descriptive statistics mean ± SEM or mean ± SD were determined for continuous variables as noted. Curves for antibody binding and neutralization were fitted after log transformation of antibody concentrations using non-linear regression analysis. Technical and biological replicates are indicated in the figure legends. Statistical analyses were performed using Prism v8.4.3 (GraphPad).

## RESOURCE AVAILABILITY

### Lead Contact

Further information and requests for resources and reagents should be directed to and will be fulfilled by the lead contact, James. E. Crowe, Jr.

### Data and Materials availability

Additional data are available in the main text or the supplementary materials. Requests for reagents may be directed to and be fulfilled by the Lead Contact: Dr. James E. Crowe, Jr. Materials reported in this study will be made available but may require execution of a Materials Transfer Agreement.

### Code availability

The computational pipeline for the clustering analysis as well as the scripts to analyze gene usages is available on GitHub: https://github.com/eccelaine/. The PyIR script used to determine sequence characteristics of each antibody is available here: https://github.com/crowelab/PyIR.

