## Supplemental Information for "Systematic analysis of human antibody response to ebolavirus glycoprotein reveals high prevalence of neutralizing public clonotypes"

Fig S1

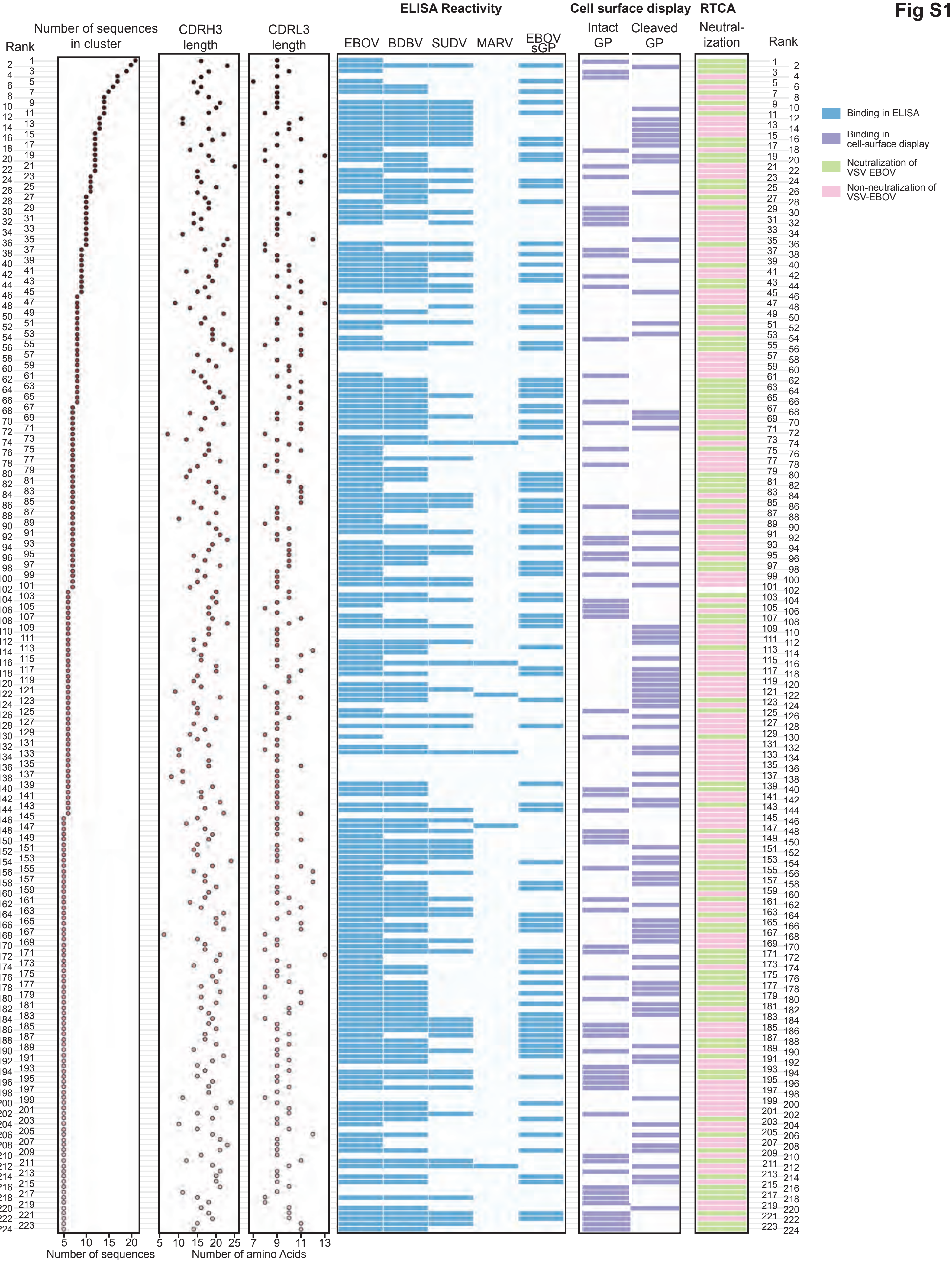
Rank
Number of sequences in cluster
CDRH3 length
CDRL3 length
ELISA Reactivity
EBOV
BDBV
SUDV
MARV
EBOV sGP
Cell surface display
Intact GP
Cleaved GP
RTCA
Neutralization
Number of sequences
Number of amino Acids
Binding in ELISA
Binding in cell-surface display
Neutralization of VSV-EBOV
Non-neutralization of VSV-EBOV

**Fig S2**

**EBOV-1182**
Heavy chain: EBOV-826
Light chain: 2.1.1D07

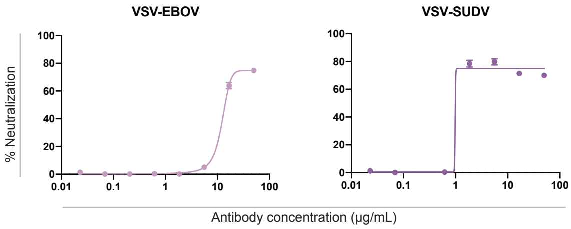


**EBOV-1190**
Heavy chain: EBOV-786
Light chain: 5.6.1A02

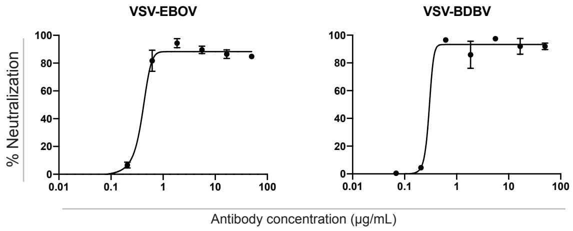


**EBOV-1187**
Heavy chain: EBOV-852
Light chain: 56-3-7A

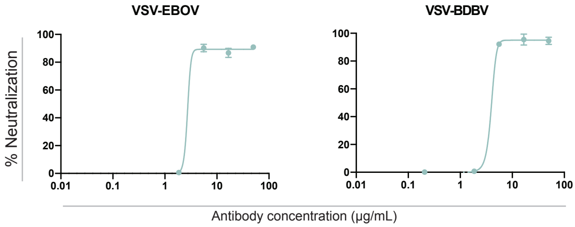

Fig S3

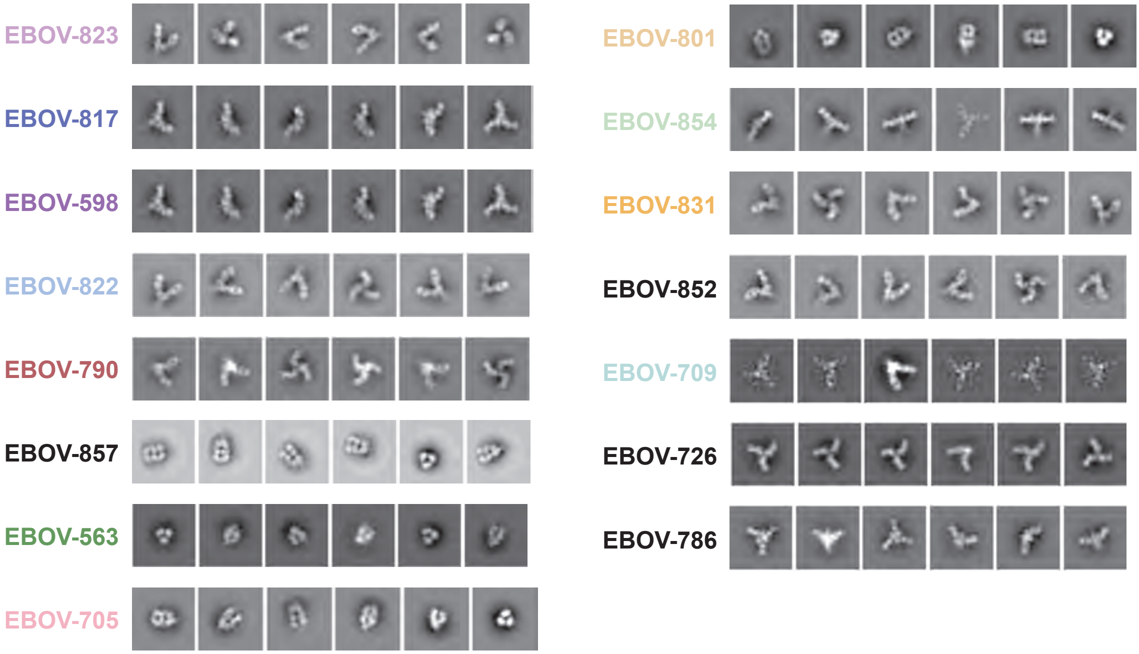
EBOV-823
EBOV-817
EBOV-598
EBOV-822
EBOV-790
EBOV-857
EBOV-563
EBOV-705
EBOV-801
EBOV-854
EBOV-831
EBOV-852
EBOV-709
EBOV-726
EBOV-786

Fig S4

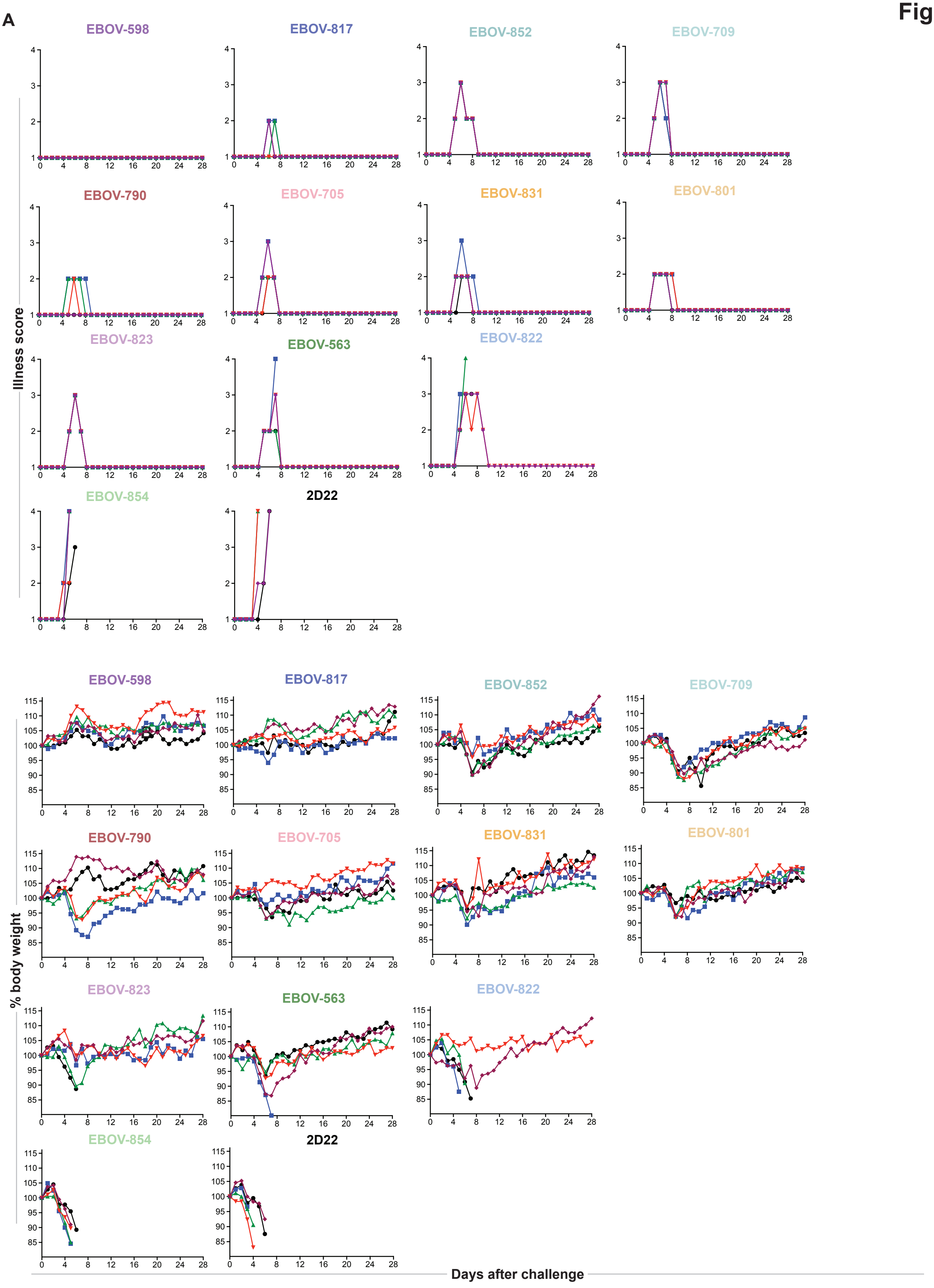
A
EBOV-598
EBOV-817
EBOV-852
EBOV-709
EBOV-790
EBOV-705
EBOV-831
EBOV-801
EBOV-823
EBOV-563
EBOV-822
EBOV-854
2D22
Illness score
% body weight
Days after challenge
